# A simple new approach to variable selection in regression, with application to genetic fine-mapping

**DOI:** 10.1101/501114

**Authors:** Gao Wang, Abhishek Sarkar, Peter Carbonetto, Matthew Stephens

## Abstract

We introduce a simple new approach to variable selection in linear regression, with a particular focus on *quantifying uncertainty in which variables should be selected*. The approach is based on a new model — the “Sum of Single Effects” (*SuSiE*) model — which comes from writing the sparse vector of regression coefficients as a sum of “single-effect” vectors, each with one non-zero element. We also introduce a corresponding new fitting procedure — Iterative Bayesian Stepwise Selection (IBSS) — which is a Bayesian analogue of stepwise selection methods. IBSS shares the computational simplicity and speed of traditional stepwise methods, but instead of selecting a single variable at each step, IBSS computes a *distribution* on variables that captures uncertainty in which variable to select. We provide a formal justification of this intuitive algorithm by showing that it optimizes a variational approximation to the posterior distribution under the *SuSiE* model. Further, this approximate posterior distribution naturally yields convenient novel summaries of uncertainty in variable selection, providing a Credible Set of variables for each selection. Our methods are particularly well-suited to settings where variables are highly correlated and detectable effects are sparse, both of which are characteristics of genetic fine-mapping applications. We demonstrate through numerical experiments that our methods outper-form existing methods for this task, and illustrate their application to fine-mapping genetic variants influencing alternative splicing in human cell-lines. We also discuss the potential and challenges for applying these methods to generic variable selection problems.

## 1 INTRODUCTION

The need to identify, or “select”, relevant variables in regression models arises in a diverse range of applications, and has spurred development of a correspondingly diverse range of methods (e.g., see O’Hara and Sillanpää, 2009; Fan and Lv, 2010; Desboulets, 2018; George and McCulloch, 1997, for reviews). However, variable selection is a complex problem, and so despite considerable work in this area there remain important issues that existing methods do not fully address. One such issue is *assessing uncertainty in which variables should be selected*, particularly in settings involving *very highly correlated variables*. Here we introduce a simple and computationally scalable approach to variable selection that helps address this issue.

Highly correlated variables pose an obvious challenge to variable selection methods, simply because they are hard to distinguish from one another. Indeed, in an extreme case where two variables (say, ***x***_1_ and ***x***_2_) are completely correlated, it is impossible to claim, based on a regression analysis, that one variable should be selected as relevant rather than the other. In some applications such ambiguity causes few practical problems. Specifically, in some applications variable selection is used only to help *build an accurate predictor*, in which case it suffices to arbitrarily select one of the two identical variables (or both); prediction accuracy is unaffected by this choice. However, in other scientific applications, variable selection is used as a means to help *learn something about the world*, and in those applications the ambiguity created by highly correlated variables is more problematic because scientific conclusions depend on which variables are selected. In these applications, it is crucial to acknowledge uncertainty in which variables should be selected. This requires methods that can draw conclusions such as “either ***x***_1_ or ***x***_2_ is relevant and we cannot decide which” rather than methods that arbitrarily select one of the variables and ignore the other. While this may seem a simple goal, in practice most existing variable selection methods do not satisfactorily address this problem (see Section 2 for further discussion). These shortcomings motivate our work here.

One particular application where these issues arise is genetic fine-mapping (e.g., Veyrieras et al., 2008; Maller et al., 2012; Spain and Barrett, 2015; Huang et al., 2017; Schaid et al., 2018). The goal of fine-mapping is to identify the genetic variants that causally affect some traits of interest (e.g., low density lipoprotein cholesterol in blood, or gene expression in cells). In other words, the main goal of fine-mapping is to learn something about the world, rather than build a better predictor. (This is not to say that predicting traits from genetic variants is not important; indeed, there is also a lot of work on prediction of genetic traits, but this is not the main goal of fine-mapping.) The most successful current approaches to fine-mapping frame the problem as a *variable selection problem*, building a regression model in which the regression outcome is the trait of interest, and the candidate predictor variables are the available genetic variants (Sillanpää and Bhattacharjee, 2005). Performing variable selection in a regression model identifies variants that may causally affect the trait. Fine-mapping is challenging because the variables (genetic variants) can be *very* highly correlated, due to a phenomenon called *linkage disequilibrium* (Ott, 1999). Indeed, typical studies contain many pairs of genetic variants with sample correlations exceeding 0.99, or even equaling 1.

Our approach builds on previous work on Bayesian variable selection in regression (BVSR) (Mitchell and Beauchamp, 1988; George and McCulloch, 1997), which has already been widely applied to genetic fine-mapping and related applications (e.g., Meuwissen et al., 2001; Sillanpää and Bhattacharjee, 2005; Servin and Stephens, 2007; Hoggart et al., 2008; Stephens and Balding, 2009; Logsdon et al., 2010; Guan and Stephens, 2011; Bottolo et al., 2011; Maller et al., 2012; Carbonetto and Stephens, 2012; Zhou et al., 2013; Hormozdiari et al., 2014; Chen et al., 2015; Wallace et al., 2015; Moser et al., 2015; Wen et al., 2016; Lee et al., 2018). BVSR is an attractive approach to these problems because it can, in principle, assess uncertainty in which variables to select, even when the variables are highly correlated. However, applying BVSR in practice remains difficult for at least two reasons. First, BVSR is computationally challenging, often requiring implementation of sophisticated Markov chain Monte Carlo or stochastic search algorithms (e.g., Bottolo and Richardson, 2010; Bottolo et al., 2011; Guan and Stephens, 2011; Wallace et al., 2015; Benner et al., 2016; Wen et al., 2016; Lee et al., 2018). Second, and perhaps more importantly, the output from BVSR methods is typically a complex posterior distribution — or samples approximating the posterior distribution — and this can be difficult to distill into results that are easily interpretable.

Our work addresses these shortcomings of BVSR through several innovations. We introduce a new formulation of BVSR, which we call the “Sum of Single Effects” *(SuSiE)* model. This model, while similar to existing BVSR models, has a different structure that naturally leads to a simple, intuitive, and fast procedure for model fitting — Iterative Bayesian Stepwise Selection (IBSS) — which is a Bayesian analogue of traditional stepwise selection methods (and which enjoys important advantages over these traditional selection methods, as we explain below). We provide a principled justification for this intuitive algorithm by showing that it optimizes a variational approximation to the posterior distribution under the *SuSiE* model. Although variational approaches to BVSR already exist (Logsdon et al., 2010; Carbonetto and Stephens, 2012), our new approach introduces a different family of approximating distribution that provides much more accurate inferences in settings with highly correlated variables.

A keyfeatureof our method, which distinguishes it from most existing BVSR methods, is that it produces “Credible Sets” of variables that quantify uncertainty in which variable should be selected when multiple, highly correlated variables compete with one another. These Credible Sets are designed to be as small as possible while still each capturing a relevant variable. Arguably, this is exactly the kind of posterior summary that one would like to obtain from MCMC-based or stochastic search BVSR methods, but doing so would require non-trivial post-processing of their output. In contrast, our method provides this posterior summary directly, and with little extra computational effort.

The structure of this paper is as follows. Section 2 provides further motivation for our work, and brief background on BVSR. Section 3 describes the new *SuSiE* model and fitting procedure. Section 4 uses simulations, designed to mimic realistic genetic fine-mapping studies, to demonstrate the effectiveness of our approach compared with existing methods. Section 5 illustrates the application of our methods to fine-mapping of genetic variants affecting splicing, and Section 6 briefly highlights the promise (and limitations) of our methods for other applications such as change-point problems. We end with a discussion highlighting avenues for further work.

## 2 BACKGROUND

### 2.1 A motivating toy example

Suppose the relationship between an *n*-vector ***y*** and an *n* × *p* matrix ***X*** = (***x***_1_,…, ***x**_p_*), is modeled as a multiple regression:

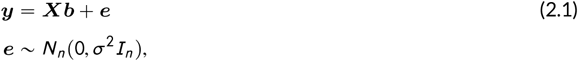

where ***b*** is a *p*-vector of regression coefficients, *e* is an *n*-vector of error terms, *σ*^2^ > 0 is the residual variance, *I_n_* is the *n* × *n* identity matrix, and *N_r_* (*μ*, Σ) denotes the *r*-variate normal distribution with mean *μ* and variance Σ. For brevity, we will refer to variables *j* with non-zero effects (*b_j_* ≠ 0) as “effect variables”.

Assume now that exactly two variables are effect variables — variables 1 and 4, say — and that these two effect variables are each completely correlated with another non-effect variable, say ***x***_1_ = ***x***_2_ and ***x***_3_ = ***x***_4_. Further suppose that no other pairs of variables are correlated. Here, because the effect variables are completely correlated with other variables, it is impossible to confidently select the correct variables, even when *n* is very large. However, given sufficient data it should be possible to conclude that there are (at least) two effect variables, and that:

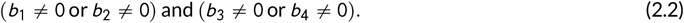

Our goal, in short, is to *provide methods that directly produce this kind of inferential statement*. Although this example is simplistic, it mimics the kind of structure that occurs in, for example, genetic fine-mapping applications, where it often happens that an association can be narrowed down to a small set of highly correlated genetic variants, but not down to an individual variant.

Most existing approaches to sparse regression do not provide statements like (2.2), nor do they attempt to do so. For example, methods that maximize a penalized likelihood, such as the lasso (Tibshirani, 1996) or elastic net (EN; Zou and Hastie, 2005), select a single “best” combination of variables, and make no attempt to assess whether other combinations are also plausible. In our toy example, EN selects all four variables, implying *b*_1_ ≠ 0, *b*_2_ ≠ 0, *b*_3_ ≠ 0 and *b*_4_ ≠ 0, which is quite different from (2.2). Recently-developed selective inference approaches (Taylor and Tibshirani, 2015) do not solve this problem, because they do not assess uncertainty in *which* variables should be selected; instead they assess uncertainty in the coefficients of the selected variables within the selected model. In our toy motivating example, selective inference methods sometimes selects the wrong variables (inevitably, due to the complete correlation with other variables) and then assigns them highly significant *p* values (see Wang et al., 2020b, for an explicit example accompanied by code). The *p* values are significant because, even though the wrong variables are selected, their coefficients — within the (wrong) selected model — can be estimated precisely. An alternative approach, which does address uncertainty in variable selection, is to control the false discovery rate (FDR) among selected variables — for example, using stability selection (Meinshausen and Bühlmann, 2010) or the knockoff filter (Barber and Candès, 2015). However, in examples with very highly correlated variables no individual variable can be confidently declared an effect variable, and so controlling the FDR among selected variables results in no discoveries, and not inferences like (2.2).

One approach to producing inferences like (2.2) is to reframe the problem, and focus on selecting *groups* of variables, rather than individual variables. A simple version of this idea might first cluster the variables into groups of highly correlated variables, and then perform some kind of “group selection” (Huang et al., 2012). However, while this could work in our toy example, in general this approach requires *ad hoc* decisions about which variables to group together, and how many groups to create — an unattractivefeature weseek to avoid. A more sophisticated version of this idea is to use hierarchical testing (Meinshausen, 2008; Yekutieli, 2008; Mandozzi and Bühlmann, 2016; Renaux et al., 2020), which requires specification of a hierarchy on the variables, but avoids an *a priori* decision on where to draw group boundaries. However, in applications where variables are not precisely arranged in a known hierarchy — which includes genetic fine-mapping — this approach is also not entirely satisfactory. In numerical assessments shown later (Section 4), we find that this approach can considerably overstate the uncertainty in which variables should be selected.

Another approach that could yield statements like (2.2), at least in principle, is the Bayesian approach to variable selection (BVSR; see Introduction for references). BVSR methods introduce a prior distribution on *b* that favours sparse models (few effect variables), and then compute a posterior distribution assessing relative support for each combination of variables. In our toy example, the posterior distribution would roughly have equal mass (≈ 0.25) on each of the four equivalent combinations {1,3}, {1,4}, {2,3} and {2,4}. This posterior distribution contains exactly the information necessary to infer (2.2). Likewise, in more complex settings, the posterior distribution contains information that could, in principle, be translated to simple statements analogous to (2.2). This translation is, however, highly non-trivial in general. Consequently, most implementations of BVSR do not provide statements like (2.2), but rather summarize the posterior distribution with a simpler but less informative quantity: the marginal posterior inclusion probability (PIP) of each variable,

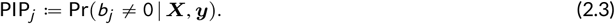

In ourexample, PIP_1_ = PIP_2_ = PIP_3_ = PIP_4_ ≈ 0.5. While not inaccurate, the PIPs do not contain the information in (2.2). In Wang et al. (2020b), we illustrate inference of Credible Sets in two additional toy examples in which the variables are correlated in more complicated ways.

### 2.2 Credible Sets

To define our main goal more formally, we introduce the concept of a *Credible Set* (CS) of variables:

#### Definition 1.

*In the context of a multiple regression model, **a level**-ρ **Credible Set** is defined to be a subset of variables that has probability ⩾ ρ of containing at least one effect variable (i.e., a variable with non-zero regression coefficient). Equivalently, the probability that all variables in the Credible Set have zero regression coefficients is* ⩽ 1 — *ρ*.

Our use of the term *Credible Set* here indicates that we have in mind a Bayesian inference approach, in which the probability statements in the definition are statements about uncertainty in which variables are selected given the available data and modelling assumptions. One could analogously define a *Confidence Set* by interpreting the probability statements as referring to the set, considered random.

Although the term *Credible Set* has been used in fine-mapping applications before, most previous uses either assumed there was a single effect variable (Malleret al., 2012), or defined a CS as a set that contains *all* effect variables (Hormozdiari et al., 2014), which is a very different definition (and, we argue, both less informative and less attainable; see further discussion below). Our definition here is closer to the “signal clusters” from Lee et al. (2018), and related to the idea of “minimal true detection” in Mandozzi and Bühlmann (2016).

With Definition 1 in place, our primary aim can be restated: we wish to report as many CSs as the data support, each with as few variables as possible. For example, to convey (2.2) we would report two CSs, {1,2} and {3,4}. As a secondary goal, we would also like to prioritize the variables within each CS, assigning each a probability that reflects the strength of the evidence for that variable being an effect variable. Our methods achieve both of these goals.

It is important to note that, if a variable is *not* included in any CS produced by our method, this does not imply that it is *not* an effect variable. This is analogous to the fact that, in hypothesis testing applications, a non-significant *p* value does not imply that the null hypothesis is true. In practice no variable selection method can guarantee identifying *every* effect variable unless it simply selects all variables, because finite data cannot rule out that every variable has a (possibly tiny) effect. This is why the CS definition of Hormozdiari et al. (2014) is unattainable, at least without strong assumptions on sparsity. It also explains why attempting to form confidence or credible sets for identifying the *true model (i.e*., the exact combination of effect variables) leads to very large sets of models; see Ferrari and Yang (2015) for example.

### 2.3 The single effect regression model

We now describe the building block for our approach, the “Single Effect Regression” (SER) model, which we define as a multiple regression model in which *exactly one of the p explanatory variables has a non-zero regression coefficient*. This idea was introduced in Servin and Stephens (2007) to fine-map genetic associations, and consequently has been adopted and extended by others, including Veyrieras et al. (2008) and Pickrell (2014). Although of very narrow applicability, the SER model is trivial to fit. Furthermore, when its assumptions hold, the SER provides exactly the inferences we desire, including CSs. For example, if we simplify our motivating example (Section 2.1) to have a single effect variable — variable 1,for example — then the SER model would, given sufficient data, infer a 95% CS containing both of the correlated variables, 1 and 2, with PIPs of approximately 0.5 each. This CS tells us that we can be confident that one of the two variables has a non-zero coefficient, but we do not know which one.

Specifically, we consider the following SER model, with hyperparameters for the residual variance, *σ*^2^, the prior variance of the non-zero effect, 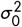, and the prior inclusion probabilities, ***π*** = (*π*_1_,…, *π_p_*), in which *π_j_* gives the prior probability that variable *j* is the effect variable:

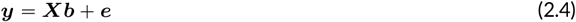

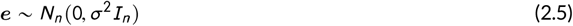

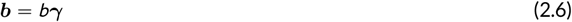

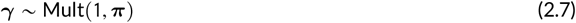

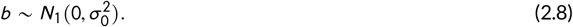

Here, ***y*** is the *n*-vector of response data; ***X*** = (***x***_1_,…, ***x***_*p*_) is an *n* × *p* matrix containing *n* observations of *p* explanatory variables; *b* is a scalar representing the “single effect”; *γ* ∈ {0,1 }^*p*^ is a *p*-vector of indicator variables; ***b*** is the *p*-vector of regression coefficients; ***e*** is an *n*-vector of independent error terms; and Mult(*m, **π***) denotes the multinomial distribution on class counts obtained when *m* samples are drawn with class probabilities given by ***π***. We assume that ***y*** and the columns of ***X*** have been centered to have mean zero, which avoids the need for an intercept term (Chipman et al., 2001).

Under the SER model (2.4–2.8), the effect vector ***b*** has exactly one non-zero element (equal to *b*), so we refer to ***b*** as a “single effect vector”. The element of ***b*** that is non-zero is determined by the binary vector ***γ***, which also has exactly one non-zero entry. The probability vector ***π*** determines the prior probability distribution on which of the *p* variables is the effect variable. In the simplest case, ***π*** = (1/*p*,…, 1/*p*); we assume this uniform prior here for simplicity, but our methods require only that ***π*** is fixed and known (so in fine-mapping one could incorporate different priors based on genetic annotations; e.g., Veyrieras et al., 2008). To lighten notation, we henceforth make conditioning on ***π*** implicit.

#### 2.3.1 Posterior under SER model

Given *σ*^2^ and 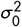, the posterior distribution on ***b*** = *γb* is easily computed:

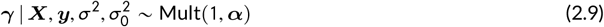

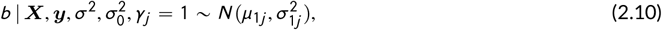

where ***α*** = (*α*_1_, …, *α_p_*) is the vector of PIPs, with 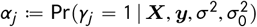, and 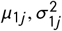 are the posterior mean and variance of *b* given *γ_j_* = 1. Calculating these quantities simply involves performing the *p* univariate regressions of ***y*** on columns *x_j_* of ***X***, for *j* = 1,…, *p*, as shown in Appendix A. From ***α***, it is also straightforward to compute a level-p CS (Definition 1), *CS*(***α***; *ρ*), as described in Maller et al. (2012), and detailed in Appendix A. In brief, this involves sorting variables by decreasing *α_j_*, then including variables in theCS until their cumulative probability exceeds *ρ*.

For later convenience, we introduce a function, *SER*, that returns the posterior distribution for ***b*** under the SER model. Since this posterior distribution is uniquely determined by the values of ***α, μ***_1_: = (*μ*_11_,…, *μ*_1*p*_) and 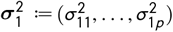 in (2.9–2.10), we can write

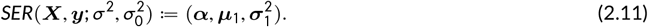

#### 2.3.2 Empirical Bayes for SER model

Although most previous treatments of the SER model assume 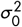 and *σ*^2^ are fixed and known, we note here the possibility of estimating 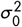 and/or *σ*^2^ by maximum-likelihood before computing the posterior distribution of ***b***. This is effectively an Empirical Bayes approach. The log-likelihood for 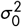 and *σ*^2^ under the SER,

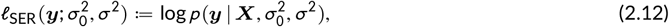

is available in closed form, and can be maximized over one or both parameters (Appendix A).

## 3 THE SUM OF SINGLE EFFECTS REGRESSION MODEL

We now introduce a new approach to variable selection in multiple regression. Our approach is motivated by the observation that the SER model provides simple inference if there is indeed exactly one effect variable; it is thus desirable to extend the SER to allow for multiple variables. The conventional approach to doing this in BVSR is to introduce a prior on ***b*** that allows for multiple non-zero entries (e.g., using a “spike-and-slab” prior; Mitchell and Beauchamp, 1988). However, this approach no longer enjoys the convenient analytic properties of the SER model; posterior distributions become difficult to compute accurately, and computing CSs is even harder.

Here we introduce a different approach which better preserves the desirable features of the SER model. The key idea is simple: introduce multiple single-effect vectors, ***b***_1_,…, ***b***_*L*_, and construct the overall effect vector, ***b***, as the sum of these single effects. We call this the “Sum of Single Effects” (*SuSiE*) regression model:

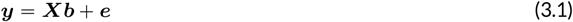

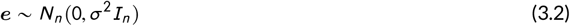

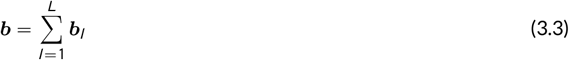

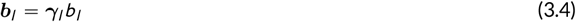

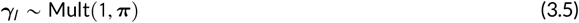

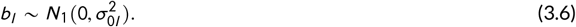

For generality, we have allowed the variance of each effect, 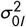, to vary among the components, *l* = 1,…, *L*. The special case in which *L* = 1 recovers the SER model. For simplicity, we initially assume *σ*^2^ and 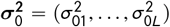 are given, and defer estimation of these hyperparameters to Section 3.1.3.

Note that if *L* ≪ *p* then the *SuSiE* model is approximately equivalent to a standard BVSR model in which *L* randomly chosen variables have non-zero coefficients (see Proposition A2 in Appendix C for a formal statement). The main difference is that with some (small) probability some of the single effects *b_l_* in the *SuSiE* model have the same non-zero co-ordinates, and so the number of non-zero elements in *b* has some (small) probability of being less than *L*. Thus, at most *L* variables have non-zero coefficients in this model. We discuss the choice of *L* in Section 3.3.

Although the *SuSiE* model is approximately equivalent to a standard BVSR model, its novel structure has two major advantages. First, it leads to a simple, iterative and deterministic algorithm for computing approximate posterior distributions. Second, it yields a simple way to calculate the CSs. In essence, because each *b_l_* captures only one effect, the posterior distribution on each *γ_l_* can be used to compute a CS that has a high probability of containing an effect variable. The remainder of this section describes both these advantages, and other issues that may arise in fitting the model.

### Algorithm 1 Iterative Bayesian stepwise selection (IBSS)

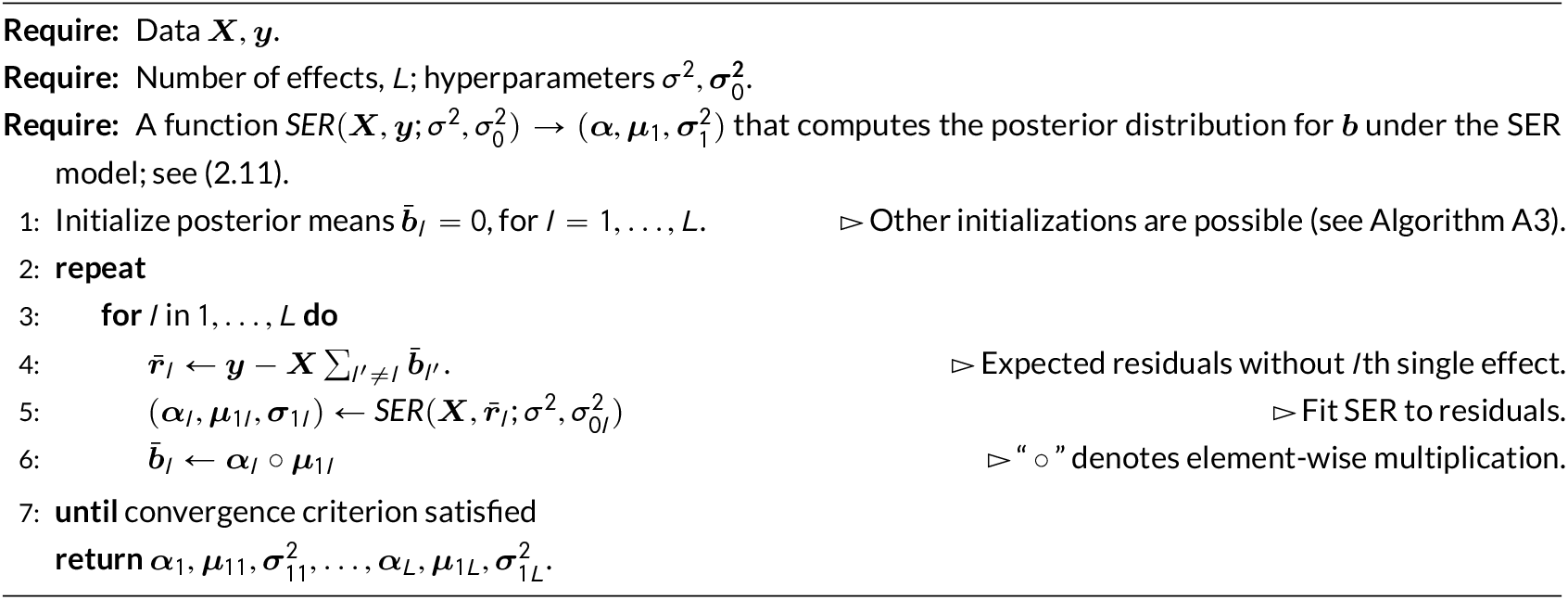

### 3.1 Fitting *SuSiE:* Iterative Bayesian stepwise selection

A key motivation for the *SuSiE* model (3.1–3.6) is that, given ***b***_1_,…, ***b***_*L*−1_, estimating ***b***_*L*_ involves simply fitting an SER model, which is analytically tractable. This immediately suggests an iterative approach to fitting this model: at each iteration use the SER model to estimate ***b**_l_* given current estimates of ***b**_l′_*, for *l*′ ≠ *l*; see Algorithm 1. This algorithm is simple and computationally scalable, with computational complexity *O*(*npL*) per outer-loop iteration.

We call Algorithm 1 “Iterative Bayesian Stepwise Selection” (IBSS) because it can be viewed as a Bayesian version of stepwise selection approaches. For example, we can compare it with an approach referred to as “forward stagewise” (FS) selection in Hastie et al. 2009, Section 3.3.3 (although subsequent literature often uses this term to mean something slightly different), also known as “matching pursuit” (Mallat and Zhang, 1993). In brief, FS first selects the single “best” variable among *p* candidates by comparing the results of the *p* univariate regressions. It then computes the residuals from the univariate regression on this selected variable, then selects the next “best” variable by comparing the results of univariate regression of the residuals on each variable. This process repeats, selecting one variable each iteration, until some stopping criterion is reached.

IBSS is similar in structure to FS, but instead of selecting a single “best” variable at each step, it computes a *distribution* on which variable to select by fitting the Bayesian SER model. Similar to FS, this distribution is based on the results of the *p* univariate regressions; consequently each selection step in IBSS has the same computational complexity as in FS, *O*(*np*). However, by computing a distribution on variables — rather than choosing a single best variable — IBSS captures uncertainty about which variable should be selected at each step. This uncertainty is taken into account when computing residuals by using a *model-averaged* (posterior mean) estimate for the regression coefficients. In IBSS, we use an iterative procedure, whereby early selections are re-evaluated in light of the later selections (as in “backfitting”; Friedman and Stuetzle, 1981). The final output of IBSS is *L* distributions on variables, parameterized by 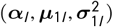, for *l* = 1,…, *L*, in place of the *L* variables selected by FS. Each distribution is easily summarized, for example, by a 95% CS for each selection.

To illustrate, consider our motivating example (Section 2.1) with ***x***_1_ = ***x***_2_, ***x***_3_ = ***x***_4_, and with variables 1 and 4 having non-zero effects. To simplify the example, suppose that the effect of variable 1 is substantially larger than the effect of variable 4. Then FS would first (arbitrarily) select either variable 1 or 2, and then select (again arbitrarily) variable 3 or 4. In contrast, given enough data, the first IBSS update would select variables 1 and 2; that is, it would assign approximately equal weights of 0.5 to variables 1 and 2, and small weights to other variables. The second IBSS update would similarly select variables 3 and 4 (again, with equal weights of approximately 0.5). Summarizing these results would yield two CSs, {1,2} and {3,4}, and the inference (2.2) is achieved. This simple example is intended only to sharpen intuition; later numerical experiments demonstrate that IBSS also works well in more realistic settings.

#### 3.1.1 IBSS computes a variational approximation to the *SuSiE* posterior distribution

The analogy between the IBSS algorithm and the simple FS procedure emphasizes the intuitive and computational simplicity of IBSS, but of course does not give it any formal support. We now provide a formal.justification for IBSS. Specifically, we show that it is a coordinate ascent algorithm for optimizing a *variational approximation (VA) to the posterior distribution* for ***b***_1_,…, ***b***_*L*_ under the *SuSiE* model (3.1–3.6). This result also suggests a method for estimating the hyperparameters *σ*^2^ and 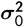.

The idea behind VA methods for Bayesian models (e.g., Jordan et al., 1999; Blei et al., 2017) is to find an approximation *q*(***b***_1_,…, ***b***_*L*_) to the posterior distribution 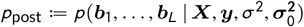 by minimizing the Kullback-Leibler (KL) divergence from *q* to *p*_post_, written as *D*_KL_(*q, p*_post_), subject to constraints on *q* that make the problem tractable. Although *D*_KL_(*q, p*_post_) itself is hard to compute, it can be formulated in terms of an easier-to-compute function, *F*, known as the “evidence lower bound” (ELBO):

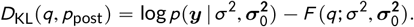

Because 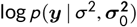 does not depend on *q*, minimizing *D*_KL_ over *q* is equivalent to maximizing *F*; and since *F* is easier to compute, this is how the problem is usually framed. See Appendix B.1 for further details. (Note that the ELBO also depends on the data, ***X*** and ***y***, but we make this dependence implicit to lighten notation.)

We seek an approximate posterior, *q*, that factorizes as

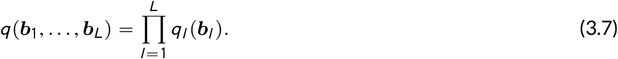

Under this approximation, ***b***_1_,…, ***b***_*L*_ are independent *a posteriori*. We make no assumptions on the form of *q_l_*; in particular, we do *not* require that each *q_l_* factorizes over the *p* elements of ***b**_l_*. This is a crucial difference from previous VA approaches for BVSR (e.g., Logsdon et al., 2010; Carbonetto and Stephens, 2012), and it means that *q_l_* can accurately capture strong dependencies among the elements of *b_l_* under the assumption that exactly one element of ***b**_l_* is non-zero. Intuitively each factor *q_l_* captures one effect variable, and provides inferences of the form that “we need one of variables {*A, B, C*}, and we are unsure about which one to select.” By extension, the approximation (3.7) provides inferences of the form “we need to select one variable among the set {*A, B, C*}, one variable among the set {*D, E, F, G*}, and so on.”

Under the assumption that the VA factorizes as (3.7), finding the optimal *q* reduces to the following problem:

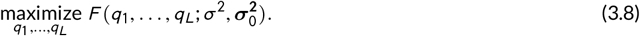

Although jointly optimizing *F* over *q*_1_,…, *q_L_* is hard, optimizing an individual factor, *q_l_*, is straightforward, and in fact reduces to fitting an SER model, as formalized in the following proposition.

##### Proposition 1.

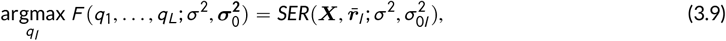

*where 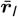 denotes the expected value of the residuals obtained by removing the estimated effects other than l*,

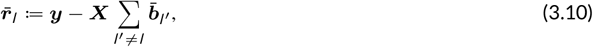

*and where* 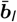 *denotes the expected value of **b**_l_ with respect to the distribution q_l_*.

For intuition, note that computing the posterior distribution for *b_l_* under (3.1 – 3.6), given the other effects *b_l′_* for *l*′ ≠ *l*, involves fitting a SER to the residuals ***y*** — ***X*** ∑_*l*′≠*l*_ ***b**_l′_*. Now consider computing an (approximate) posterior distribution for ***b**_l_* when ***b**_l’_* are not known, and we have approximations *q_l_*’ to their posterior distributions. Proposition 1 states that we can solve for argmax*_q_l__ F*(*q*_1_,…, *q_L_*) using a similar procedure, except that each *b_l_*’ is replaced with the (approximate) posterior mean 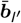.

The following is an immediate consequence of Proposition 1:

##### Corollary 1.

*IBSS (Algorithm 1) is a coordinate ascent algorithm for maximizing the ELBO, F, over q satisfying* (3.7). *Equivalently, it is a coordinate ascent algorithm for minimizing the KL divergence D*_KL_(*q, p*_post_) *over q satisfying* (3.7), *where p*_post_ *is the true posterior distribution under the SuSiE model*.

Further, as a consequence of being a coordinate ascent algorithm, IBSS converges to a stationary point of *F* under conditions that are easily satisfied:

##### Proposition 2.

*Provided that 0* < ***σ, σ***_0_ < ∞ *and **π**_j_* > 0 *for all j* = 1,…, *p, the sequence of iterates q generated by the IBSS method (parameterized by* 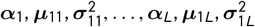) *converges to a limit point that is a stationary point of F*.

The proof of Propositions 1 and 2 and Corollary 1 are given in Appendix B.

#### 3.1.2 Contrast to previous variational approximations

A critical point is that the VA being computed by IBSS is different from previous “fully factorized” VAs for BVSR (e.g., Logsdon et al., 2010; Carbonetto and Stephens, 2012). In settings with highly correlated variables, the new VA produces results that are not only *quantitatively* different, but also *qualitatively* different from the fully factorized VA. For example, in our motivating example (Section 2.1), the new VA provides statements like (2.2), whereas the fully factorized VAs do not. Rather, a fully factorized VA often selects at most one of two identical variables without adequately capturing uncertainty in which variable should be selected (Carbonetto and Stephens, 2012). This feature makes the fully factorized VA unsuitable for applications where it is important to assess uncertainty in *which variables are selected*.

More generally, the new VA computed by IBSS satisfies the following intuitive condition: when two variables are identical, inferences drawn about their coefficients are identical (assuming the priors on their coefficients are the same). Despite the simplicity of this condition, it is not satisfied by existing VAs, nor by point estimates from penalized likelihood approaches with *L*_0_ or *L*_1_ penalty terms. (In fact, Zou and Hastie 2005 use this condition as motivation for the elastic net method, which does ensure that point estimates for coefficients of identical variables are equal.) This property is formalized in the following proposition.

##### Proposition 3.

*Consider applying the IBSS algorithm (Algorithm 1) to a data set in which two columns of **X** are identical; that is, **x**_j_ = **x**_k_ for some j ≠ k. Further suppose that the prior distributions on selecting these two variables are equal (π_j_ = π_k_). Then the approximate posterior computed by IBSS will be exchangeable in j, k; that is, if ω_jk_*: ∝^*p*^ → ∝^*p*^ *denotes the function that permutes elements j and k of a p-vector, and q denotes the approximate posterior obtained from the IBSS algorithm, then*

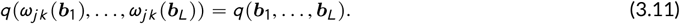

*Proof*. Since 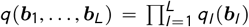, it suffices to show that each *q_l_* is exchangeable in *j, k; i.e., q_l_* (*ω_jk_* (*b_l_*)) = *q_l_*(*b_l_*) for all *l* = 1,…, *L*. This exchangeability is satisfied after every iteration of the IBSS algorithm because the algorithm computes *q_l_* (parameterized by 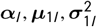) as the exact posterior distribution under an SER model (Step 5 of Algorithm 1), and this posterior is exchangeable in *j, k* because both the prior and likelihood are exchangeable.

Because the exchangeability is satisfied after every iteration of IBSS, and not j’ust at convergence, the result is not sensitive to stopping criteria. By contrast, the corresponding EN property (Zou and Hastie, 2005) holds only at convergence — for example, in numerical implementations of the EN method (e.g., the `glmnet` R package), the coefficient estimates for identical variables can differ substantially. Similarly, MCMC-based implementations of BVSR may satisfy this exchangeability property only asymptotically.

#### 3.1.3 Estimating *σ*^2^, 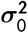

Algorithm 1 can be extended to estimate the hyperparameters *σ*^2^ and 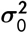 by adding steps to maximize 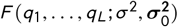 over *σ*^2^ and/or 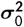. Estimating the hyperparameters by maximizing the ELBO can be viewed as an EM algorithm (Dempster et al., 1977) in which the E-step is approximate (Heskes et al., 2004; Neal and Hinton, 1998).

Optimizing *F* over *σ*^2^ involves computing the expected residual sum of squares under the VA, which is straightforward; see Appendix B for details.

Optimizing *F* over 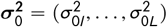 can be achieved by modifying the step that computes the posterior distribution for *b_l_* under the SER model to first estimate the hyperparameter 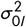 in the SER model by maximum likelihood; that is, by maximizing the *SER* likelihood (2.12) over 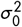, keeping *σ*^2^ fixed (Step 5 of Algorithm A3). This is a one-dimensional optimization which is easily performed numerically (we used the R function `optim`).

Algorithm A3 in Appendix B extends Algorithm 1 to include both these steps.

### 3.2 Posterior inference: posterior inclusion probabilities and Credible Sets

Algorithm 1 provides an approximation to the posterior distribution of ***b*** under the *SuSiE* model, parameterized by 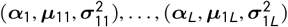. From these results it is straightforward to compute approximations to various posterior quantities of interest, including PIPs and CSs.

#### 3.2.1 Posterior inclusion probabilities

Under the *SuSiE* model, the effect of explanatory variable *j* is 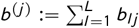, which is zero if and only if *b_lj_* = 0 for all *l* = 1,…, *L*. Under our VA the *b_lj_* are independent across *l*, and therefore

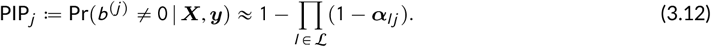

Here, we set 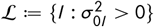 to treat the case where some 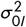 are zero, which can happen if 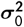 is estimated.

#### 3.2.2 Credible Sets

Computing the sets *CS*(***α***_*l*_; *ρ*) (A.4),for *l* = 1,…, *L* immediately yields *L* CSs that satisfy Definition 1 under the VA to the posterior.

If *L* exceeds the number of detectable effects in the data, then in practice many of the *L* CSs are large, often containing the majority of variables. The intuition is that once all the detectable signals have been accounted for, the IBSS algorithm becomes very uncertain about which variable to include at each step, and so the distributions *α_l_* become very diffuse. CSs that contain very many uncorrelated variables are of essentially no inferential value — whether or not they contain an effect variable — and so in practice it makes sense to ignore them. To automate this, in this paper we discard CSs with “purity” less than 0.5, where we define purity as the smallest absolute correlation among all pairs of variables within the CS. (To reduce computation for CSs containing over 100 variables, we sampled 100 variables at random to estimate the purity.) The purity threshold of 0.5 was chosen primarily for comparing with Lee et al. (2018), who use a similar threshold in a related context. While any choice of threshold is somewhat arbitrary, in practice we observed that most CSs are either very pure (> 0.95) or very impure (< 0.05), with intermediate cases being rare (Figure S2),so most results are robust to this choice of threshold.

### 3.3 Choice of *L*

It may seem that *SuSiE* would be sensitive to the choice of *L*. In practice, however, key inferences are often robust to overstating *L*; for example, in our simulations below, the simulated number of effects was between 1 and 5, whereas we still obtain good results with *L* = 10. This is because, when *L* is larger than necessary, the method is very uncertain about where to place the extra effects — consequently, it distributes them broadly among many variables, and therefore they are too diffuse to impact key inferences. For example, setting *L* to be larger than necessary inflates the PIPs of many variables, but only slightly, and the extra components result in CSs with low purity.

While inferences are generally robust to overstating *L*, we also note that the Empirical Bayes version of our method, which estimates 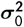, also effectively estimates the number of effects: when *L* is greater than the number of signals in the data, the maximum likelihood estimate of 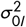 will be zero or close to zero for many *L*, which in turn forces *b_l_* to zero. This is closely related to the idea behind “automatic relevance determination” (Neal, 1996).

### 3.4 Identifiability and label-switching

The parameter vectors ***b***_1_,…, ***b**_L_* introduced in the *SuSiE* model are technically non-identifiable, in that the likelihood *p*(***y***|***X***, *σ*^2^, ***b***_1_,…, ***b***_*L*_) is unchanged by permutation of the labels 1,…, *L*. As a result, the posterior distribution *p*_post_ is symmetric with respect to permutations of the labels (assuming the prior is also symmetric) — that is, for any permutation, *v*: {1,…, *L*} → {1,…, *L*}, we have *p*(***b***_1_,…, ***b***_*L*_ | ***X***, ***y***, *σ*^2^) = *p*(***b***_*v*(1)_,…, ***b***_*v*(*L*)_ | ***X, y***, *σ*^2^). A similar non-identifiability also occurs in mixture models, where it is known as the “label-switching problem” (Stephens, 2000).

In principle, non-identifiability due to label-switching does not complicate Bayesian inference; the posterior distribution is well-defined, and correctly reflects uncertainty in the parameters. In practice, however, complications can arise. Specifically, the label-switching typically causes the posterior distribution to be multi-modal, with *L*! symmetric modes corresponding to the *L*! different labellings (*v* above). Care is then needed when summarizing this posterior distribution. For example, the posterior mean will not be a sensible estimate for ***b***_1_,…, ***b***_*L*_ because it averages over the *L*! modes (Stephens, 2000).

Fortunately, our variational approximation (Section 3.1.1) avoids these potential complications of label-switching. This is due to the way that variational approximations behave when approximating the posterior distribution of a mixture model; they typically produce a good approximation to one of the permutations, effectively ignoring the others (Wang and Titterington, 2006; Blei et al., 2017; Pati et al., 2018). See also the discussion of “spontaneous symmetry-breaking” in Wainwright and Jordan (2007). Consequently, our posterior approximation *q*(***b***_1_,…, ***b***_*L*_) approximates just one of the L! symmetric modes of the true posterior, avoiding the issues with label-switching that can occur when summarizing the true posterior distribution.

Formally this non-identifiability causes the objective *F* optimized by the IBSS algorithm to be invariant to relabeling — that is,

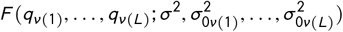

is the same for all permutations *ν* — and therefore every solution 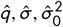 returned by our IBSS algorithm has *L*! equivalent solutions that achieve the same value of the ELBO, *F*, each corresponding to a different labeling (and a different mode of the true posterior). These *L*! solutions are inferentially equivalent; they all imply the same distribution for the unordered set {***b***_1_,…, ***b***_*L*_}, the same distribution for the sum 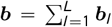 (which does not depend on the labeling), and they all produce the same PIPs and the same CSs. Thus, it does not matter which mode is used.

## 4 NUMERICAL COMPARISONS

We performed numerical comparisons on data generated to closely mimic our main motivating application: genetic fine-mapping. Specifically, we generated data for fine-mapping of expression quantitative trait loci (eQTLs), which are genetic variants associated with gene expression. We used these simulations to assess our methods, and compare with state-of-the-art BVSR methods that were specifically developed for this problem. We also compared against a (frequentist) hierarchical testing method (Mandozzi and Bühlmann, 2016; Renaux et al., 2020).

In genetic fine-mapping, ***X*** is a matrix of genotype data, in which each row corresponds to an individual, and each column corresponds to a genetic variant, typically a single nucleotide polymorphism (SNP). In our simulations, we used the real human genotype data from n = 574 genotype samples collected as part of the Genotype-Tissue Expression (GTEx) proj’ect (GTEx Consortium, 2017). To simulate fine-mapping of locally-acting variants associated with gene expression (*cis* eQTLs), we randomly selected 150 genes out of the > 20,000 genes on chromosomes 1-22, then assigned ***X*** to be the genotypes for genetic variants nearby the transcribed region of the selected gene. For a given gene, between *p* = 1,000 and *p* = 12,000SNPs were included in the fine-mapping analysis; for more details on how SNPs were selected, see Appendix D.

These real genotype matrices, **X**, exhibit complex patterns of correlations; see Figure A1 for example. Furthermore, many variables are strongly correlated with other variables: fora randomly chosen variable, the median number of other variables with which its correlation exceeds 0.9 is 8; and the median number of other variables with which its correlation exceeds 0.99 is 1. Corresponding means are even larger — 26 and 8 other variables, respectively — because some variables are strongly correlated with hundreds of other variables. Thus these genotype matrices lead to challenging, but realistic, variable selection problems.

We generated synthetic outcomes ***y*** under the multiple regression model (2.1), with assumptions on ***b*** specified by two parameters: 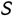, the number of effect variables; and *ϕ*, the proportion of variance in *y* explained by ***X*** (abbreviated as “PVE”). Given 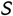 and *ϕ*, we simulated ***b*** and ***y*** as follows:

1. Sample the indices of the 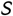 effect variables, 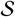, uniformly at random from {1,…, *p*}.
2. For each 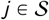, independently draw *b_j_* ~ *N*(0,0.6^2^), and for all 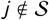, set *b_j_* = 0.
3. Set *σ*^2^ to achieve the desired PVE, *ϕ*; specifically wesolvefor *σ*^2^ in 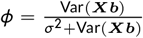, where Var(·) denotessample variance.
4. For each *i* = 1,…, *n*, draw 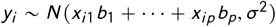.

We generated data sets under two simulation scenarios. In the first scenario, each data set has *p* = 1,000 SNPs. We generated data sets using all pairwise combinations of 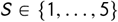 and *ϕ* ∈ {0.05,0.1,0.2,0.4}. These settings were chosen to span typical expected values for eQTL studies. We simulated two replicates for each gene and for each combination of 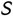 and *ϕ*. Therefore, in total we generated 2 × 150 × 5 × 4 = 6,000 data sets for the first simulation scenario.

In the second simulation scenario, we generated data sets with all *cis*SNPs (defined as SNPs within 1 Megabase radius from transcription start site of a gene), ranging from 3,000 to 12,000 SNPs, and to generate the outcomes ***y***, we set 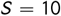 and *ϕ* = 0.3. We generated two replicates for each gene, resulting in a total of 2 × 150 = 300 data sets in the second simulation scenario.

### 4.1 Illustrative example

We begin with an example to illustrate that the IBSS algorithm (Algorithm 1) can perform well in a challenging fine-mapping setting. This example is summarized in Figure 1.

**FIGURE 1.**
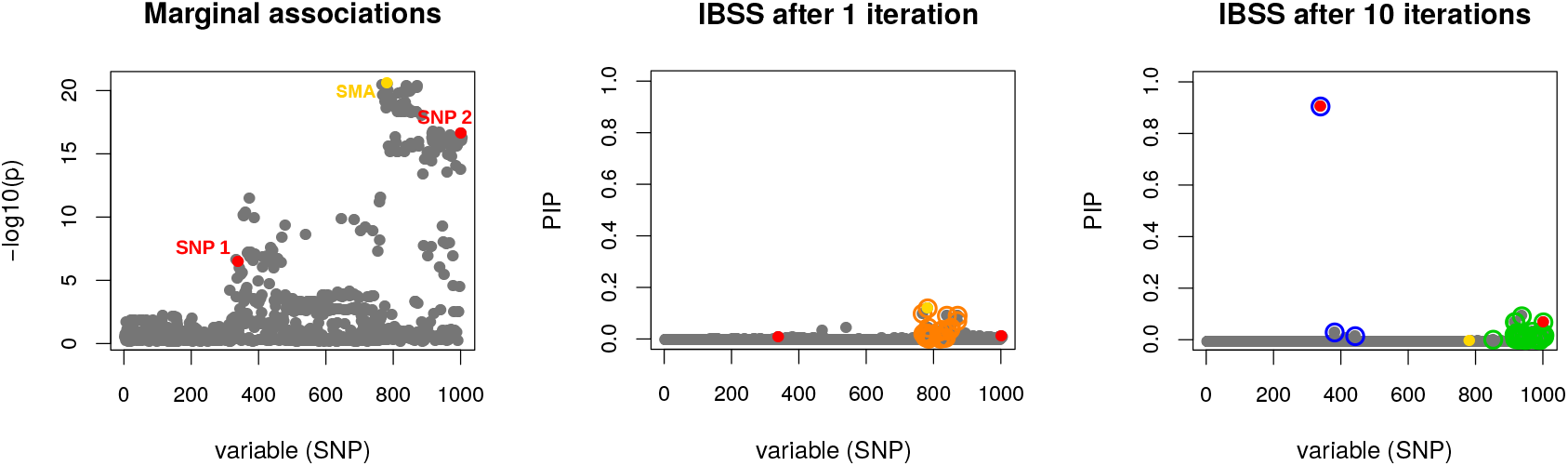
Fine-mapping example to illustrate that IBSS algorithm can deal with a challenging case. Results are from a simulated data set with *p* = 1,000 variables (SNPs). Someof these variables are very strongly correlated (Figure A1). Two out of the 1,000 variables are effect variables (red points, labeled “SNP1” and “SNP 2” in the left-hand panel). We chose this example from our simulations because the strongest marginal association (SMA) is a non-effect variable (yellow point, labeled “SMA” in the left-hand panel). After 1 iteration (middle panel), IBSS incorrectly identifies a CS containing the SMA and no effect variable (orange points). However, after 10 iterations (and also at convergence) the IBSS algorithm has corrected itself (right-panel), finding two 95% CSs — dark blue and green open circles — each containing a true effect variable. Additionally, neither CS contains the SMA variable. One CS (blue) contains only 3 SNPs (purity of 0.85), whereas the other CS (green) contains 37 very highly correlated variables (purity of 0.97). In the latter CS, the individual PIPs are small, but the inclusion of the 37 variables in this CS indicates, correctly, high confidence in at least one effect variable among them.

We draw this example from one of our simulations in which the variable with the strongest marginal association (SMA) with ***y*** is not one of the actual effect variables (in this example, there are two effect variables). This situation occurs because the SMA variable has moderate correlation with both effect variables, and these effects combine to make its marginal association stronger than the marginal associations of the individual effect variables. Standard forward selection in this case would select the wrong (SMA) variable in the first step; indeed, after one iteration, IBSS also yields a CS that includes the SMA variable (Figure 1, middle panel). However, as the IBSS algorithm proceeds, it recognizes that, once other variables are accounted for, the SMA variable is no longer required. After 10 iterations (at which point the IBSS solution is close to convergence) IBSS yields two high-purity CSs, neither containing the SMA, and each containing one of the effect variables (Figure 1, right panel). Our manuscript resource repository includes an animation showing the iteration-by-iteration progress of the IBSS algorithm (Wang et al., 2020a).

This example, where the SMA variable does not appear in a CS, also illustrates that multiple regression can sometimes result in very different conclusions than a marginal association analysis.

### 4.2 Posterior inclusion probabilities

Next, we seek to assess the effectiveness of our methods more quantitatively. We focus initially on one of the simpler tasks in BVSR: computing posterior inclusion probabilities (PIPs). Most implementations of BVSR compute PIPs, making it possible to compare results across several implementations. Here we compare our methods (henceforth *SuSiE*, implemented in R package susieR, version 0.4.29) with three other software implementations specifically developed for genetic fine-mapping applications: CAVIAR (Hormozdiari et al., 2014, version 2.2), FINEMAP (Benner et al., 2016, version 1.1) and DAP-G (Wen et al., 2016; Lee et al., 2018, installed using source code from the git repository, commit id `efllb26`). These methods are all implemented as C++ programs. They implement similar BVSR models, and differ in the algorithms used to fit these models and the priors on the effect sizes. CAVIAR exhaustively evaluates all possible combinations of up to *L* non-zero effects among the *p* variables. FINEMAP and DAP-G approximate this exhaustive approach by heuristics that target the best combinations. Another important difference among methods is that FINEMAP and CAVIAR perform inference using summary statistics computed from each data set — specifically, the marginal association *Z* scores and the *p × p* correlation matrix for all variables — whereas, as we apply them here, DAP-G and *SuSiE* use the full data. The summary statistic approach can be viewed as approximating inferences from the full data; see Lee et al. (2018) for discussion.

For *SuSiE*, we set *L* = 10 for all data sets generated in the first simulation scenario, and *L* = 20 for the second scenario. We assessed performance when both estimating the hyperparameters 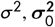, and when fixing one or both of these hyperparameters. Overall performance of these different approaches were similar, and here we show results when *σ*^2^ was estimated, and when 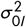 was fixed to 0.1 Var(***y***) (consistent with data applications in Section 5); other results are in Supplementary Data (Figure S4 and Figure S5). Parametersettingsfor other methods are given in Appendix D. We ran CAVIAR and FINEMAP only on simulations with 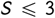 since these methods are computationally more intensive than the others (particularly for larger 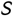).

Since these methods differ in their modelling assumptions, one should not expect their PIPs to be equal. Nonetheless, we found generally reasonably good agreement (Figure 2A). For 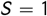, the PIPs from all four methods agree closely. For 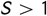, the PIPs from different methods are also highly correlated; correlations between PIPs from *SuSiE* and other methods vary from 0.94 to 1 across individual data sets, and the number of PIPs differing by more than 0.1 is always small — the proportions vary from 0.013% to 0.2%. In the scatterplots, this agreement appears less strong because the eye is drawn to the small proportion of points that lie away from the diagonal, but the vast majority of points lie on or nearthe origin. In addition, all four methods produce reasonably well-calibrated PIPs (Figure S1).

**FIGURE 2.**
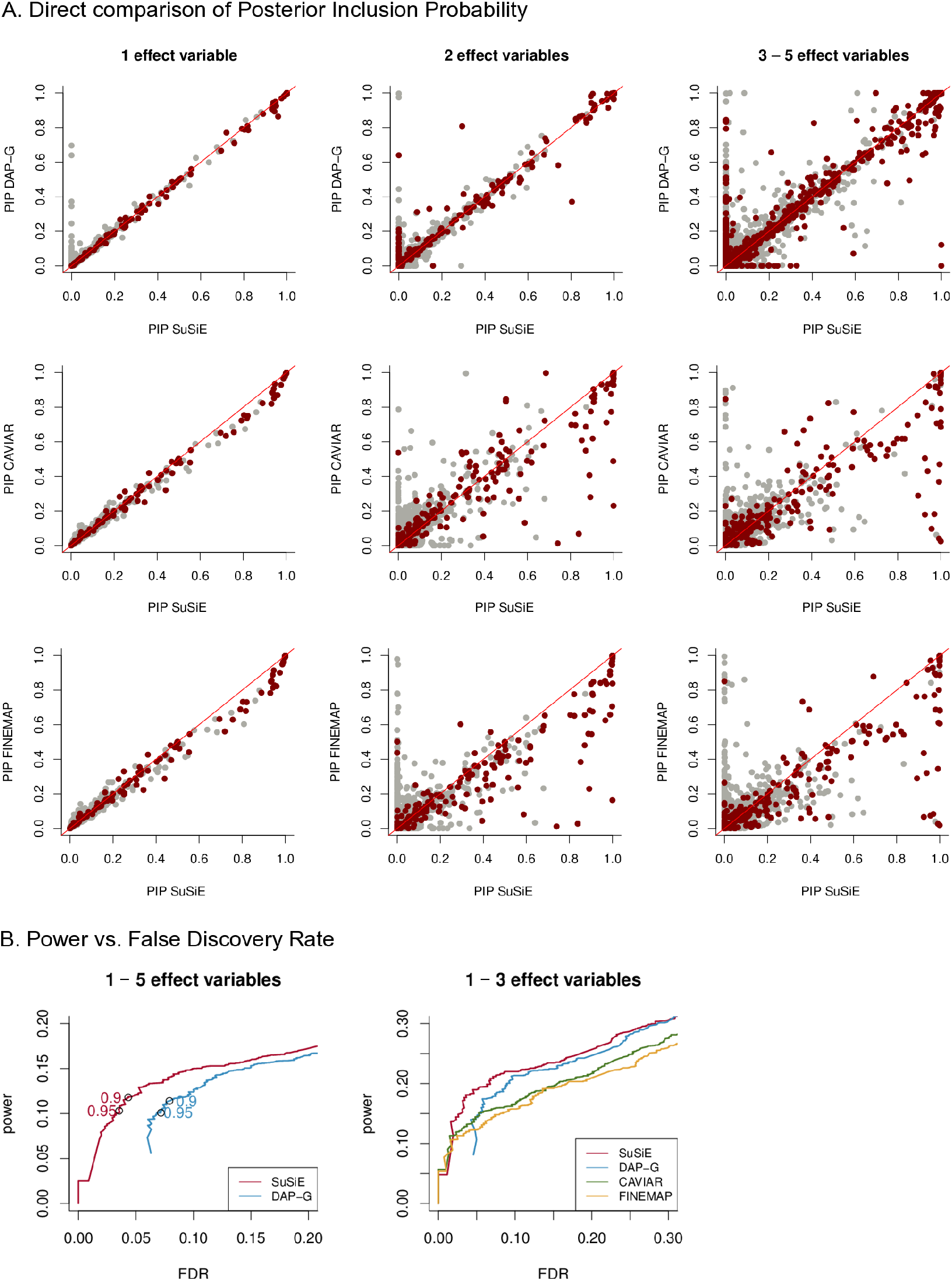
Evaluation of posterior inclusion probabilities (PIPs). Scatterplots in **Panel A** compare PIPs computed by *SuSiE* against PIPs computed using other methods (DAP-G, CAVIAR, FINEMAP). Each point depicts a single variable in one of the simulations: dark red points represent true effect variables, whereas light gray points represent variables with no effect. The scatterplot in Panel B combine results across the first set of simulations. **Panel B** summarizes power versus FDR from the first simulation scenario of. These curves are obtained by independently varying the PIP threshold for each method. The open circles in the left-hand plot highlight power versus FDR at PIP thresholds of 0.9 and 0.95). These quantities are calculated as 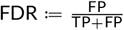 (also known as the “false discovery proportion”) and 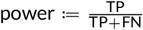, where FP,TP, FN and TN denote the number of False Positives, True Positives, False Negatives and True Negatives, respectively. (This plot is the same as a *precision-recall curve* after reversing the x-axis, because 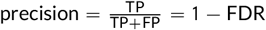, and recall = power.) Note that CAVIAR and FINEMAP were run only on data sets with 1 — 3 effect variables.

The general agreement of PIPs from four different methods suggests that: (i) all four methods are mostly accurate for computing PIPs for the data set sizes explored in our numerical comparisons; and (ii) the PIPs themselves are usually robust to details of the modelling assumptions. Nonetheless, some non-trivial differences in PIPs are clearly visible from Figure 2A. Visual inspection of these differences suggests that the *SuSiE* PIPs may better distinguish effect variables from non-effect variables, in that there appears a higher ratio of red-gray points below the diagonal than above the diagonal. This is confirmed in our analysis of power versus False Discovery Rate (FDR), obtained by varying the PIP threshold independently for each method; at a given FDR, the *SuSiE* PIPs always yield higher power (Figure 2B).

Notably, even though *SuSiE* is implemented in R, its computations are much faster than the other methods implemented in C++: for example, in the data sets simulated with 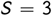, *SuSiE* is, on average, roughly 4 times faster than DAP-G, 30 times faster than FINEMAP, and 4,000 times faster than CAVIAR (Table 1).

**TABLE 1:** Runtimes from data sets simulated with 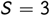 (all runtimes are in seconds)

| method | mean | min. | max. |
| --- | --- | --- | --- |
| SuSiE | 0.64 | 0.34 | 2.28 |
| DAP-G | 2.87 | 2.23 | 8.87 |
| FINEMAP | 23.01 | 10.99 | 48.16 |
| CAVIAR | 2907.51 | 2637.34 | 3018.52 |

Because *SuSiE* computations scale linearly with data size (computational complexity *O*(*npL*) per iteration) it can easily handle data sets much larger than the ones in these simulations. To illustrate, running *SuSiE* (*L* = 10) on two larger simulated data sets — one with *n* = 100, 000, *p* = 500; another with *n* = 1, 000, *p* = 50, 000; each with 4 effect variables — took 25s and 43s on a modern Linux workstation (see Appendix D.2 for details). This is competitive with lasso, implemented in the `glmnet` R package, version 2.0.18, which with 10-fold cross-validation (and other parameters at their defaults) took 82s for each data set.

In summary, in the settings considered here,*SuSiE* produces PIPs that are as or more reliable than existing BVSR methods, and does so at a fraction of the computational effort.

### 4.3 Credible Sets

#### Comparison with DAP-G

A key feature of *SuSiE* is that it yields multiple Credible Sets (CSs), each aimed at capturing an effect variable (Definition 1). The only other BVSR method that attempts something similar, as far as we are aware, is DAP-G, which outputs signal clusters defined by heuristic rules (Lee et al., 2018). Although the authors do not refer to their signal clusters as CSs, and they do not give a formal definition of signal cluster, the intent of these signal clusters is similar to our CSs, and so for brevity we henceforth refer to them as CSs.

We compared the level 95% CSs produced by *SuSiE* and DAP-G in several ways. First we assessed their empirical (frequentist) coverage levels; that is, the proportion of CSs that contain an effect variable. Since our CSs are Bayesian Credible Sets, 95% CSs are not designed, or guaranteed, to have frequentist coverage of 0.95 (Fraser, 2011). Indeed, coverage will inevitably depend on simulation scenario; for example, in completely null simulations, in which the data are simulated with ***b*** = **0**, *every* CS would necessarily contain no effect variable, and so the coverage would be zero Nonetheless, under reasonable circumstances that include effect variables, one might hope that the Bayesian CSs would have coverage near the nominal levels. And, indeed, we confirmed this was the case: in the simulations, CSs from both methods typically had coverage slightly below 0.95, and in most cases above 0.90 (Figure 3; see Figure S3 for additional results).

**FIGURE 3.**
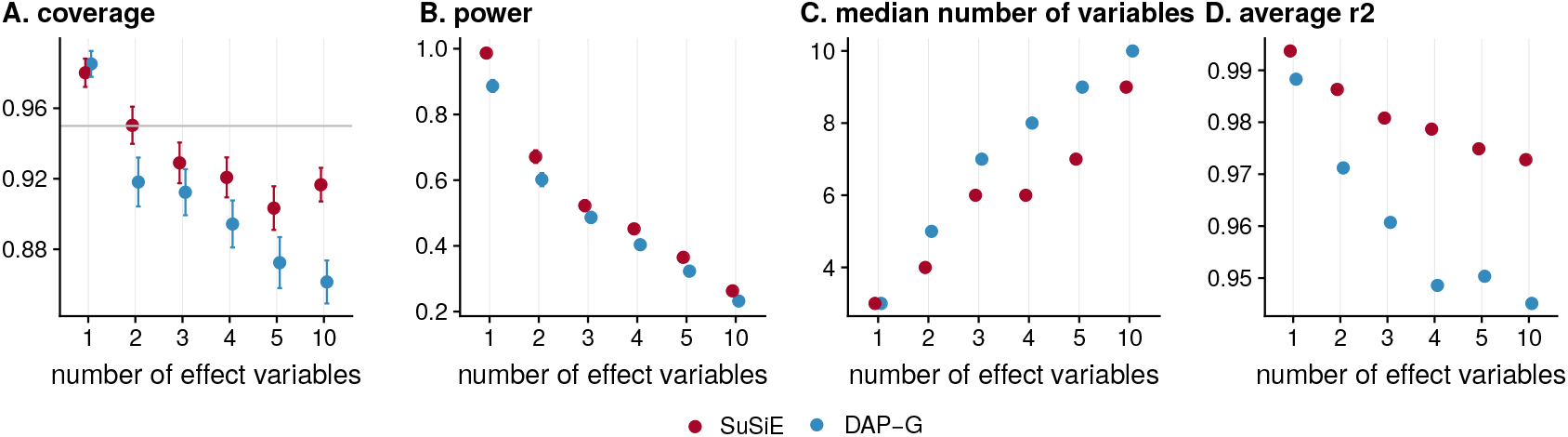
Comparison of 95% credible sets (CS) from *SuSiE* and DAP-G. Panels show A) coverage, B) power, C) median size and D) average squared correlation of the variables in each CS. These statistics are taken as mean over all CSs computed in all data sets; error bars in Panel A show 2 × standard error. Simulations with 1-5 effect variables are from the first simulation scenario, and simulations with 10 effect variables are from the second scenario.

Having established that the methods produce CSs with similar coverage, we compared them by three other criteria: (i) power (overall proportion of simulated effect variables included in a CS); (ii) average size (median number of variables included in a CS) and (iii) purity (here, measured as the average squared correlation of variables in a CS since this statistic is provided by DAP-G). By all three metrics, the CSs from *SuSiE* are consistently an improvement over DAP-G—they achieve higher power, smaller size, and higher purity (Figure 3).

Although the way that we construct CSs in *SuSiE* does not require that they be disjoint, we note that the CSs rarely overlapped (after filtering out low purity CSs; see Section 3.2.2). Indeed, across the thousands of simulations, there was only one example of two CSs overlapping.

#### Comparison with hierarchical testing

Finally, we compared our CSs with results produced by the R package `hierinf` (Renaux et al., 2020) (version 1.3.1), which implements a frequentist approach to identifying significant clusters of variables based on hierarchical testing (Meinshausen, 2008; Mandozzi and Bühlmann, 2016). In brief, this approach starts by assuming that the variables are organized in a given hierarchy. Then, starting from the top of the hierarchy, it proceeds to test whether groups of variables (clades in the hierarchy) contain at least one non-zero effect. Each time a group is deemed significant, the method proceeds to test clades in the next level of the hierarchy. The procedure ultimately reports the smallest significant clades detected, where the significance criteria are designed to control the overall family-wise error rate (FWER) at a pre-specified level, *α*. We note that FWER control is not guaranteed when *p > n* and variables are highly correlated (Mandozzi and Bühlmann, 2016), which is the situation in our simulations.

Although the theory for controlling FWER in hierarchical testing is elegant, genetic variants do not come in a natural hierarchy, and so for fine-mapping the need to specify a hierarchy is a drawback. Here we use the `cluster _var` function from `hierinf`, which infers a hierarchical clustering. There is no simple correspondence between the level α and (frequentist) coverage rates of the significant clusters, so selecting a suitable α is non-trivial; in our simulations, we found that empirical coverage was typically close to 0.95 when α = 0.1, so we report results for α = 0.1.

The results (Table 2) show that the `hierinf` clusters are substantially larger, and have lower purity than the CSs from *SuSiE*, as well as DAP-G. For example, in simulations with 5 effect variables, the *SuSiE* CSs have a median size of 7 variables with an average *r*^2^ of 0.97, whereas the `hierinf` clusters have a median size of 54 variables with an average *r*^2^ of 0.56. Further, *SuSiE* and DAP-G achieved greater power — that is, they identified more credible sets containing true signals — than the significant clusters from `hierinf`.

**TABLE 2:** Comparison of CSs from *SuSiE* and DAP-G to significant clusters from hierarchical inference (`hierinf` software, with FWER level *α* = 0.1). Results are averages across all data sets in the first simulation scenario.

| #effects | power | | | coverage | | | median size | | | average $r^2$ | | |
| --- | --- | --- | --- | --- | --- | --- | --- | --- | --- | --- | --- | --- |
|  | SuSiE | DAP-G | hierinf | SuSiE | DAP-G | hierinf | SuSiE | DAP-G | hierinf | SuSiE | DAP-G | hierinf |
| 1 | 0.99 | 0.89 | 0.97 | 0.98 | 0.99 | 0.94 | 3 | 3 | 8 | 0.99 | 0.99 | 0.82 |
| 2 | 0.67 | 0.60 | 0.55 | 0.95 | 0.92 | 0.96 | 4 | 5 | 20 | 0.99 | 0.97 | 0.71 |
| 3 | 0.52 | 0.49 | 0.39 | 0.93 | 0.91 | 0.95 | 6 | 7 | 34 | 0.98 | 0.96 | 0.64 |
| 4 | 0.45 | 0.40 | 0.29 | 0.92 | 0.89 | 0.95 | 6 | 8 | 37 | 0.98 | 0.95 | 0.60 |
| 5 | 0.37 | 0.32 | 0.24 | 0.90 | 0.87 | 0.98 | 7 | 9 | 54 | 0.97 | 0.95 | 0.56 |

We believe that the much larger number of variables included in the `hierinf` clusters partly reflects a fundamental limitation of the hierarchical approach to this problem. Specifically, by assuming a hierarchy that does not truly exist, the method artificially limits the clusters of variables it can report. This will sometimes force it to report clusters that are larger than necessary. For example, with 3 variables, if variables 2 and 3 are grouped together at the bottom of the hierarchy, then the method could never report a cluster {1,2}, representing the statement “either variable 1 or 2 is an effect variable, but we cannot tell which,” even if the data support such an inference. Instead, it would have to report the larger cluster, {1,2,3}.

While our work here was under peer review, and available as a preprint (Wang et al., 2019), we became aware of new related work in a preprint by Sesia et al. (2020). Similar to `hierinf` this new method tests groups of variables at multiple resolutions in a hierarchy; but it improves on `hierinf` by controlling the false discovery rate of selected groups (rather than type I error), and with statistical guarantees that hold even in the presence of highly correlated variables. Comparisons with our method find that their significant groups are typically larger than ours (Sesia et al., 2020, Figure 4), presumably in part due to the fundamental limitation with the hierarchical approach (discussed above).

**FIGURE 4.**
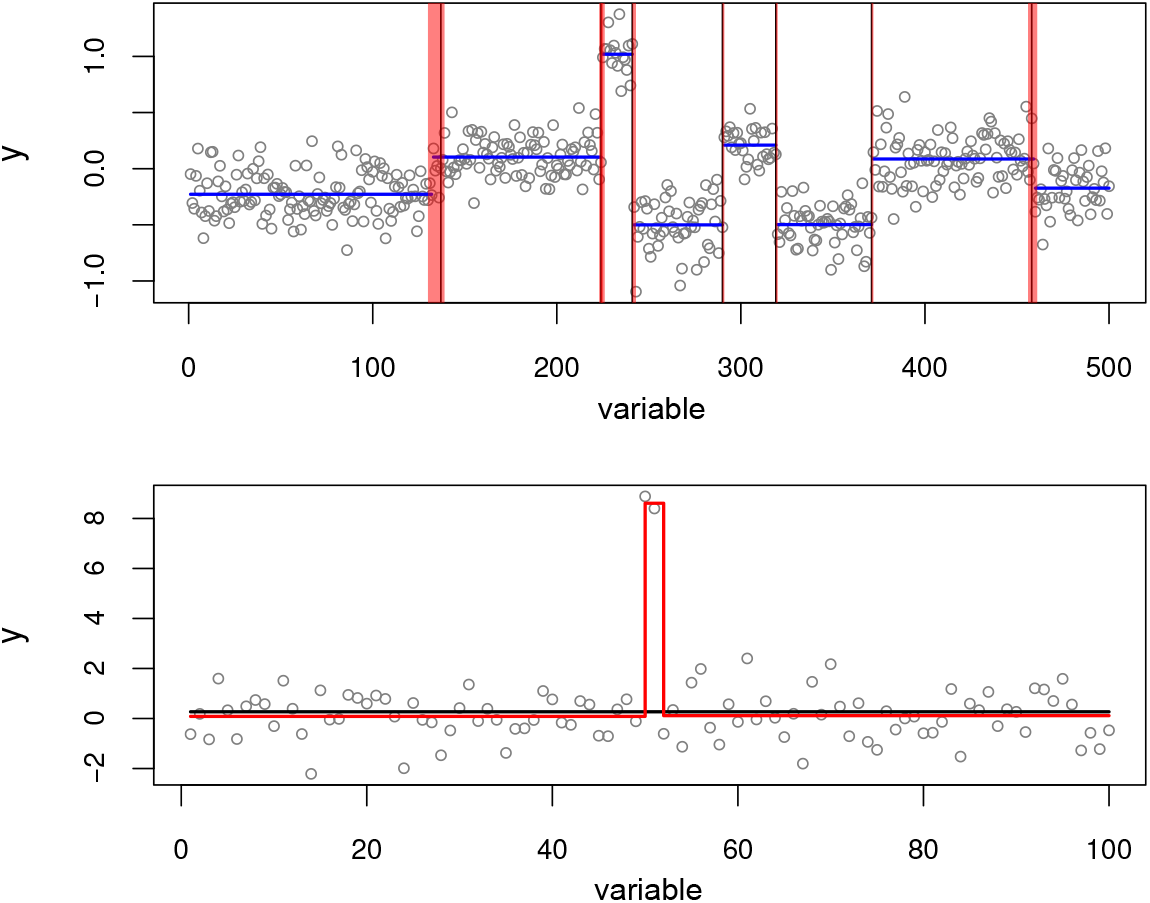
Illustration of *SuSiE* applied to two change point problems. The **top panel** shows a simulated example with seven change points (the vertical black lines). The blue horizontal lines show the mean function inferred by the `segment` method from the `DNAcopy` R package (version 1.56.0). The inference is reasonably accurate — all change points except the left-most one are nearly exactly recovered — but provides no indication of uncertainty in the locations of the change points. The red regions depict the 95% CSs for change point locations inferred by *SuSiE;* in this example, every CS contains a true change point. The **bottom panel** shows a simulated example with two change points in quick succession. This example is intended to illustrate convergence of the IBSS algorithm to a (poor) local optimum. The black line shows the fit from the IBSS algorithm when it is initialized to a null model in which there are no change points; this fit results in no change points being detected. The red line also shows the result of running IBSS, but this time the fitting algorithm is initialized to the true model with two change points. The latter accurately recovers both change points, and attains a higher value of the objective function (—148.2 versus —181.8).

## 5 APPLICATION TO FINE-MAPPING SPLICING QTLS

To illustrate *SuSiE* for a real fine-mapping problem, we analyzed data from Li et al. (2016) aimed at detecting genetic variants (SNPs) that influence splicing (known as “splicing QTLs”, sQTLs). These authors quantified alternative splicing by estimating, at each intron in each sample, a ratio capturing how often the intron is used relative to other introns in the same “cluster” (roughly, gene). The data involve 77,345 intron ratios measured on lymphoblastoid cell lines from 87 Yoruban individuals, together with genotypes of these individuals. Following Li et al. (2016), we preprocessed the intron ratios by regressing out the first 3 principle components of the matrix of intron ratios; the intent is to control for unmeasured confounders (Leek and Storey, 2007). For each intron ratio, we fine-mapped SNPs within 100 kb of the intron, which is approximately 600 SNPs on average. In short, we ran *SuSiE* on 77,345 data sets with *n* = 87 and *p* ≈ 600.

To specify the prior variance 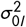, we first estimated typical effect sizes from the data on all introns. Specifically, we performed univariate (SNP-by-SNP) regression analysis at every intron, and estimated the PVE of the top (strongest associated) SNP. The mean PVE of the top SNP across all introns was 0.096, so we applied *SuSiE* with 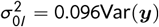, and with the columns of ***X*** standardized to have unit variance. The residual variance parameter *σ*^2^ was estimated by IBSS.

We then ran *SuSiE* to fine-map sQTLs at all 77,345 introns. After filtering for purity, this yielded a total of 2,652 CSs (level 0.95) spread across 2,496 intron units. These numbers are broadly in line with the original study, which reported 2,893 significant introns at 10% FDR. Of the 2,652 CSs identified, 457 contain exactly one SNP, representing strong candidates for being the causal variants that affect splicing. Another 239 CSs contain exactly two SNPs. The median size of a CS was 7, and the median purity was 0.94.

The vast majority of intron units with a CS had exactly one CS (2,357 of 2,496). Thus, *SuSiE* could detect at most one sQTLfor most introns. Of the remainder, 129 introns yielded 2 CSs, 5 introns yielded 3 CSs, 3 introns yielded 4 CSs, and 2 introns yielded 5 CSs. This represents a total of 129 + 10 + 9 + 8 = 156 additional (“secondary”) signals that would be missed in conventional analyses that report only one signal per intron. Both primary and secondary signals were enriched in regulatory regions (Appendix E), lending some independent support that *SuSiE* is detecting real signals. Although these data show relatively few secondary signals, this is a small study (*n* = 87); in larger studies, the ability of *SuSiE* to detect secondary signals will likely be greater.

## 6 AN EXAMPLE BEYOND FINE-MAPPING: CHANGE POINT DETECTION

Although our methods were motivated by genetic fine-mapping, they are also applicable to other sparse regression problems. Here we apply *SuSiE* to an example quite different from fine-mapping: change point detection. This application also demonstrates that the IBSS algorithm can sometimes produce a poor fit — due to getting stuck in a local optimum — which was seldom observed in our fine-mapping simulations. We believe that examples where algorithms fail are just as important as examples where they succeed — perhaps more so — and that this example could motivate improvements.

We consider a simple change point model

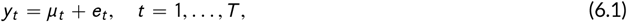

where *t* indexes a dimension such as space or time, and the errors *e_t_* are independently normal with zero mean and variance *σ*^2^. The mean vector ***μ*** — (*μ*_1_,…, ***μ**_T_*) is assumed to be piecewise constant; the indices *t* where changes to ***μ*** occur, *μ_t_* ≠ *μ*_*t* +1_, are called the “change points.”

To capture change points being rare, we formulate the change point model as a sparse multiple regression (2.1) in which ***X*** has *T* — 1 columns, and the tth column is a step function with a step at location *t*; that is, ***x***_*st*_ = 0 for *s ≤ t*, and ***x***_*st*_ = 1 for all *s > t*). The tth element of ***b*** then determines the change in the mean at position *t, μ*_*t* +1_ — *μ_t_*. Therefore, the non-zero regression coefficients in this multiple regression model correspond to change points in ***μ***.

The design matrix ***X*** in this setting has a very special structure, and quite different from fine-mapping applications; the (*T* — 1) × (*T* — 1) correlation matrix decays systematically and very slowly away from the diagonal. By exploiting this special structure of ***X***, *SuSiE* computations can be made *O*(*TL*) rather than the *O*(*T*^2^ *L*) of a naive implementation; for example, the matrix-vector product ***X***^*T*^***y***, naively an *O*(*T*^2^) computation, can be computed as the cumulative sum of the elements of the reverse of ***y***, which is an *O*(*T*) computation.

Change point detection has a wide range of potential applications, such as segmentation of genomes into regions with different numbers of copies of the genome. Software packages in R that can be used for detecting change points include `changepoint` (Killick and Eckley, 2014), `DNAcopy` (Seshan and Olshen, 2018; Olshen et al., 2004), `bcp` (Erdman and Emerson, 2007) and `genlasso` (Tibshirani, 2014; Arnold and Tibshirani, 2016); see Killick and Eckley (2014) for a longer list. Of these, only `bcp`, which implements a Bayesian method, quantifies uncertainty in estimated change point locations, and bcp provides only PIPs, not CSs for change point locations. Therefore, the ability of *SuSiE* to provide CSs is unusual, and perhaps unique, among existing change point detection methods.

To illustrate its potential for change point estimation, we applied *SuSiE* to a simulated example included with the `DNAcopy` R package. In this example, all settings for running *SuSiE* remain unchanged from the fine-mapping simulations (Section 4). The top label of Figure 4 shows results of applying *SuSiE* and `DNAcopy` to the data set. Both methods provide accurate estimates of the change points; indeed all change point locations except the left-most one are recovered nearly exactly. However, only *SuSiE* provides 95% CSs for each estimate of a change point location. And, indeed, *SuSiE* is most uncertain about the left-most change point. All the true change points in this example are contained in a *SuSiE* CS, and every CS contains a true change point. This occurs even though we set *L* = 10 to be greater than the number of true change points (7); the three extra CSs were filtered out because they contained variables that were very uncorrelated. (To be precise, *SuSiE* reported 8 CSs after filtering, but two of the CSs overlapped and contained the same change point; this observation of overlapping of CSs contrasts with the fine-mapping simulations in Section 4 where overlapping CSs occurred very rarely.)

To demonstrate that IBSS can converge to a poor local optimum, consider the simulated example shown in the bottom panel of Figure 4, which consists of two change points in quick succession that cancel each other out (the means before and after the change points are the same). This example was created specifically to illustrate a limitation of the IBSS procedure: IBSS can only introduce or update one change point at a time, and every update is guaranteed to increase the objective, whereas in this example introducing one change point will make the fit worse. Consequently, when *SuSiE* is run from a null initialization, IBSS finds no change points, and reports no CSs.

This poor outcome represents a limitation of the IBSS algorithm, not a limitation of the *SuSiE* model or the variational approximation. To show this, we re-ran the IBSS algorithm, but initializing at a solution that contained the two true change points. This yielded a fit with two CSs, each containing the one of the correct change points. This also resulted in a much improved value of the objective function (—148.2 versus —181.8). Better algorithms for fitting *SuSiE* models, or more careful initializations of IBSS, will be needed to address this shortcoming,

## 7 DISCUSSION

We have presented a simple new approach to variable selection in regression. Compared with existing methods, the main benefits of our approach are its computational efficiency, and its ability to provide CSs summarizing uncertainty in which variables should be selected. Our numerical comparisons demonstrate that for genetic fine-mapping our methods outperform existing methods at a fraction of the computational cost.

Although our methods apply generally to variable selection in linear regression, further work may be required to improve performance in difficult settings. In particular, while the IBSS algorithm worked well in our fine-mapping experiments, for change point problems we showed that IBSS may converge to poor local optima. We have also seen convergence problems in experiments with many effect variables (e.g. 200 effect variables out of 1,000). Such problems may be alleviated by better initialization, for example using fits from convex objective functions (e.g., lasso) or from more sophisticated algorithms for non-convex problems (Bertsimas et al., 2016; Hazimeh and Mazumder, 2018). More ambitiously, one could attempt to develop better algorithms to reliably optimize the *SuSiE* variational objective function in difficult cases. For example, taking smaller steps each iteration, rather than full coordinate ascent, may help.

At its core, the *SuSiE* model is based on adding up simple models (SERs) to create more flexible models (sparse multiple regression). This additive structure is the key to our variational approximations, and indeed our methods apply generally to adding up any simple models for which exact Bayesian calculations are tractable, not only SER models (Appendix B; Algorithm A2). These observations suggest connections with both additive models and boosting (e.g., Friedman et al., 2000; Freund et al., 2017). However, our methods differ from most work on boosting in that each “weak learner” (here, SER model) itself yields a model-averaged predictor. Other differences include our use of backfitting, the potential to estimate hyper-parameters by maximizing an objective function rather than cross-validation, and the interpretation of our algorithm as a variational approximation to a Bayesian posterior. Although we did not focus on prediction accuracy here, the generally good predictive performance of methods based on model averaging and boosting suggest that *SuSiE* should work well for prediction as well as variable selection.

It would be natural to extend our methods to generalized linear models (GLMs), particularly logistic regression. In genetic studies with small effects, Gaussian models are often adequate to model binary outcomes (e.g. Pirinen et al., 2013; Zhou et al., 2013). However, in other settings this extension may be more important. One strategy would be to directly modify the IBSS algorithm, replacing the SER fitting procedure with a logistic or GLM equivalent. This strategy is appealing in its simplicity, although it is not obvious what objective function the resulting algorithm is optimizing. Alternatively, for logistic regression one could use the variational approximations developed by Jaakkola and Jordan (2000).

For genetic fine-mapping, it would also be useful to modify our methods to deal with settings where only summary data are available (e.g. the *p* univariate regression results). Many recent fine-mapping methods deal with this (e.g., Chen et al., 2015; Benner et al., 2016; Newcombe et al., 2016) and ideas used by these methods can also be applied to *SuSiE*. Indeed, our software already includes preliminary implementations for this problem.

Beyond genetic fine-mapping, one could consider applying *SuSiE* to related tasks, such as genetic prediction of complex traits and heritability estimation (Yang et al., 2011). However, we do not expect *SuSiE* to provide substantial improvements over existing methods for these tasks. This is because, in general, the best existing approaches to these problems do not make strict sparsity assumptions on the effect variables; they allow for models in which many (or all) genetic variants affect the outcome (Meuwissen et al., 2001; Moser et al., 2015; Speed and Balding, 2014; Vilhjálmsson et al., 2015; Zhou et al., 2013). Nonetheless, it is possible that the ideas introduced here for sparse modelling could be combined with existing methods allowing non-sparse effects to improve prediction and heritability estimation, similar to Zhou et al. (2013).

Finally, we are particularly interested in extending these methods to select variables simultaneously for multiple outcomes *(multivariate regression* and *multi-task learning)*. Joint analysis of multiple outcomes should greatly enhance power and precision to identify relevant variables (e.g., Stephens, 2013). The computational simplicity of our approach makes it particularly appealing for this complex task, and we are currently pursuing this direction by combining our methods with those from Urbut et al. (2019).

## 8 DATA AND RESOURCES

*SuSiE* is implemented in the R package susieR available at https://github.com/stephenslab/susieR. Source code and a website detailing the analysis steps for numerical comparisons and data applications are available at our manuscript resource repository (Wang et al., 2020b), also available at https://github.com/stephenslab/susie-paper.

## Acknowledgements

Wethank Kaiqian Zhang and Yuxin Zou for their substantial contributions to the development and testing of the susieR package. Computing resources were provided by the University of Chicago Research Computing Center. This work was supported by NIH grant HG002585 and by a grant from the Gordon and Betty Moore Foundation (grant GBMF #4559).

## Appendices

## A DETAILS OF POSTERIOR COMPUTATIONS FOR THE SER MODEL

### A.1 Bayesian simple linear regression

To derive posterior computations for the SER model (2.4–2.8), it helps to start with an even simpler (univariate) linear regression model:

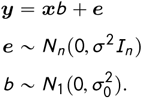

Here, ***y*** is an *n*-vector of response data (centered to have mean zero), ***x*** is an *n*-vector containing values of a single explanatory variable (similarly centered), ***e*** is an *n*-vector of independent error terms with variance *σ*^2^, *b* is the scalar regression coefficient to be estimated, 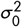 is the prior variance of *b*, and *I_n_* is the *n* × *n* identity matrix.

Given *σ*^2^ and 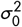, the posterior computations for this model are very simple; they can be conveniently written in terms of the usual least-squares estimate of *b*, 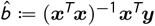, its variance 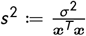, and the corresponding *z* score, 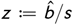. The posterior distribution for *b* is where

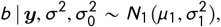

where

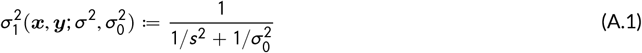

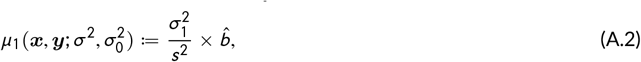

and the Bayes Factor (BF) for comparing this model with the null model (*b* = 0) is

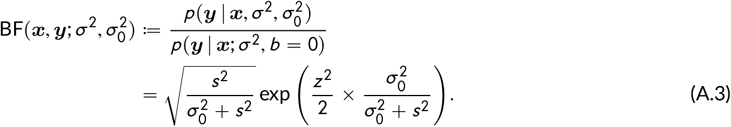

This expression matches the “asymptotic BF” of Wakefield (2009), but here, because we consider linear regression given *σ*^2^, it is an exact expression for the BF, not just asymptotic.

### A.2 The single effect regression model

Under the SER model (2.4–2.8), the posterior distribution of (*b*_1_, …, *b_p_*) = (*bγ*_1_,…, *bγ_p_*) conditioned on 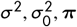 is given in the main text (eqs. 2.9 and 2.10), and is reproduced here for convenience:

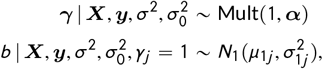

where the vector of posterior inclusion probabilities (PIPs), ***α*** = (*α*_1_,…, *α_p_*), can be expressed in terms of the simple linear regression BFs (A.3),

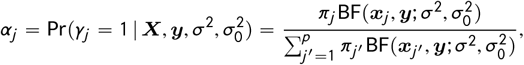

where *μ*_1*j*_ and 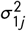. are the posterior mean (A.2) and variance (A.1) from the simple regression model of ***y*** on ***x***_*j*_:

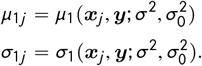

Our algorithm requires the first and second moments of this posterior distribution, which are

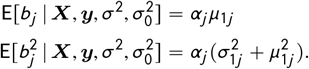

### A.3 Computing Credible Sets

As noted in the main text, under the SER model it is straightforward to compute a level-*ρ* CS (Definition 1), *CS*(***α***; *ρ*). The procedure is given in Maller et al. (2012), and for convenience we describe it here as well.

Given ***α***, let *r* = (*r*_1_,…, *r_p_*) denote the indices of the variables ranked in order of decreasing *α_j_*, so that 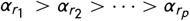, and let 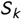 denote the cumulative sum of the *k* largest PIPs:

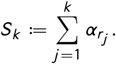

Now take

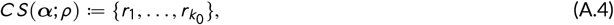

where 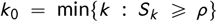. This choice of *k*_0_ ensures that the CS is as small as possible while satisfying the requirement that it is a level-*ρ* CS.

### A.4 Estimating hyperparameters

As noted in the main text, it is possible to take an empirical Bayes approach to estimating the hyperparameters 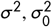. The likelihood is

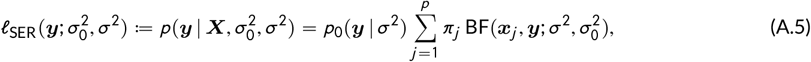

where *p*_0_ denotes the distribution of ***y*** under the “null” that *b* = 0 (i.e. 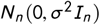), and 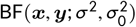 is given in eq. A.3. The likelihood (A.5) can be maximized over one or both parameters using available numerical algorithms.

## B DERIVATION OF VARIATIONAL ALGORITHMS

### B.1 Background: Empirical Bayes and variational approximation

Here we introduce some notation and elementary results which are later applied to our specific application.

#### B.1.1 Empirical Bayes as a single optimization problem

Consider the following generic model:

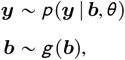

where ***y*** represents a vector of observed data, ***b*** represents a vector of unobserved (latent) variables of interest, 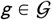 represents a prior distribution for **b** (which in the empirical Bayes paradigm is treated as an unknown to be estimated), and *θ* ∈ Θ represents an additional set of parameters to be estimated. This formulation also includes as a special case situations where ***g*** is pre-specified rather than estimated simply by making 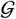 contain a single distribution.

Fitting this model by empirical Bayes typically involves the following two steps:

1. Obtain estimates 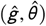 of (*g, θ*) by maximizing the log-likelihood:

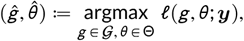

where

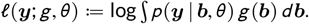
2. Given these estimates, 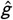 and 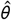, compute the posterior distribution for ***b***,

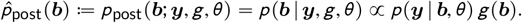 This two-step procedure can be conveniently expressed as a single optimization problem:

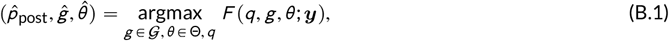

with

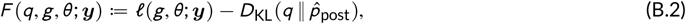

and where

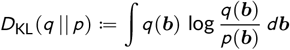

is the Kullback-Leibler (KL) divergence from *q* to *p*, and the optimization of *q* in (B.1) is over *all possible distributions on **b***. The function *F* (B.2) is often called the “evidence lower bound”, or ELBO, because it is a lower bound for the “evidence” (the marginal log-likelihood). (This follows from the fact that KL divergence is always non-negative.)

This optimization problem (B.1) is equivalent to the usual two-step EB procedure. This equivalence follows from two observations:

1. Since the marginal log-likelihood, *ℓ*, does not depend on *q*, we have

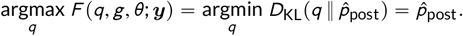
2. Since the minimum of *D*_KL_ with respect to *q* is zero for any (*θ, g*), we have that max_*q*_ *F*(*q, g, θ; **y***) = *ℓ*(***y**; g, θ*),and as a result

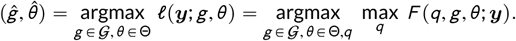

#### B.1.2 Variational approximation

The optimization problem (B.1) is often intractable. The idea of variational approximation is to adjust the problem to make it tractable, simply by restricting the optimization over all possible distributions on ***b*** to 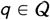, where 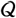 denotes a suitably chosen class of distributions. Therefore, we seek to solve B.1 subject to the additional constraint that 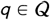:

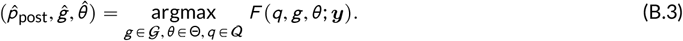

From the definition of *F*, it follows that optimizing *F* over 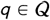 (for a given *g* and *θ*) corresponds to minimizing the KL divergence from *q* to the posterior distribution, and so can be interpreted as finding the “best” approximation to the posterior distribution for ***b*** among distributions in the class 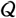. And the optimization of *F* over (*g, θ*) can be thought of as replacing the optimization of the log-likelihood with optimization of a lower bound to the log-likelihood (the ELBO).

We refer to solutions of the general problem (B.1), in which *q* is unrestricted, as “empirical Bayes (EB) solutions,” and we refer to solutions of the restricted problem (B.3) as “variational empirical Bayes (VEB) solutions.”

#### B.1.3 Form of ELBO

It is helpful to note that, by simple algebraic manipulations, the ELBO (B.2) can be decomposed as

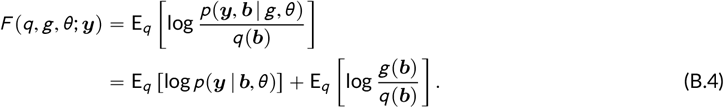

### B.2 The additive effects model

We now apply the above results to fitting an additive model, 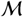, that includes the SuSiE model (3.1–3.6) as a special case:

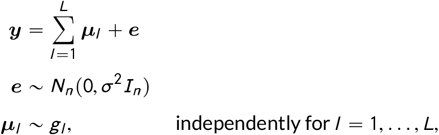

where 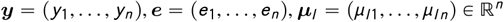. We let 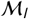 denote the simpler model that is derived from 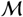 by setting *μ_l′_* = 0 for all *l′* ≠ *l* (*i.e*., 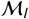 is the model that includes only the *l*th additive term), and we use *ℓ_l_* to denote the marginal log-likelihood for this simpler model:

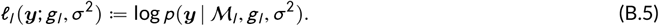

The SuSiE model corresponds to the special case of 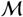 where ***μ***_*l*_ = ***Xb***_*l*_,for *l* = 1,…, *L*, and each *g_l_* is the “single effect prior” in (2.6–2.8). Further, in this special case each 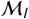 is a “single effect regression” (SER) model (2.4–2.8).

The key idea introduced in this section is that we can fit 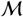 by variational empirical Bayes (VEB) provided we can fit each simpler model 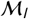 by EB. To expand on this, consider fitting the model 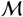 by VEB, where the restricted family 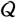 is the class of distributions on (***μ***_1_,…, ***μ***_*L*_) thatfactorize over ***μ***_1_,…, ***μ***_*L*_; that is, for any 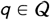,

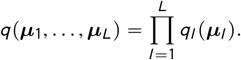

For 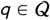, using expression (B.4), we obtain the following expression for the ELBO, *F*:

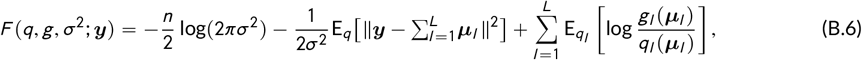

in which ||·|| denotes the Euclidean norm, and *g* denotes the collection of priors (*g*_1_,…, *g_L_*). The expected value in the second term of (B.6) is the expected residual sum of squares (ERSS) under the variational approximation *q*, and depends on *q* only through its first and second moments. Indeed, if we denote the posterior first and second moments by

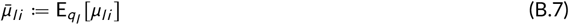

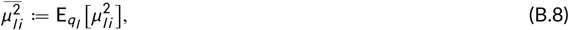

and we define 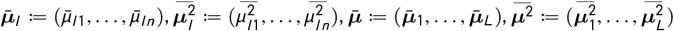, then we have that

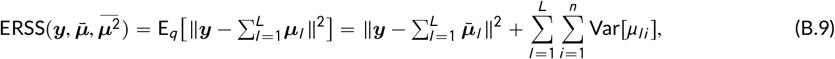

where 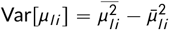. This expression follows from the definition of the expected residual sum of squares, and from independence across *l* = 1,…, *L*, after some algebraic manipulation; see Section B.7.

Fitting 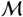 by VEB involves optimizing *F* in (B.6) over *q, g, σ*^2^. Our strategy is to update each (*q_l_, g_l_*) for *l* = 1,…, *L* while keeping *σ*^2^ and other elements of q, g fixed, and with a separate optimization step for *σ*^2^ with *q, g* fixed. This strategy is summarized in Algorithm A1.

#### Algorithm A1 Coordinate ascent for fitting additive model ℳ by VEB (outline)

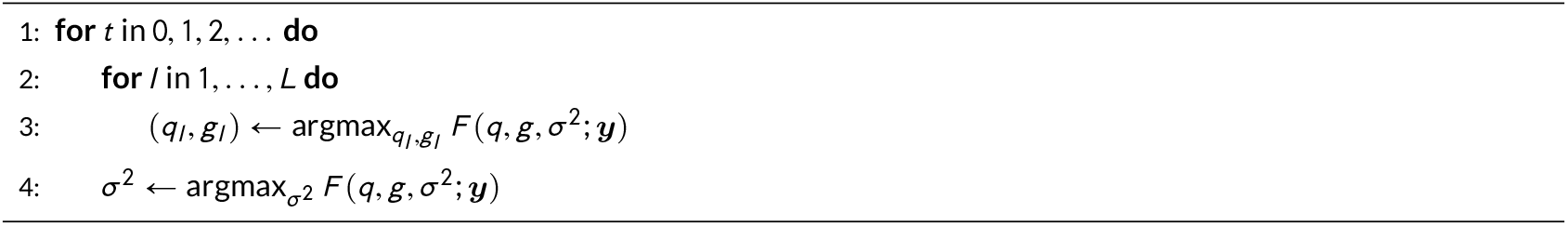

#### Algorithm A2 Coordinate ascent for fitting additive model ℳ by VEB

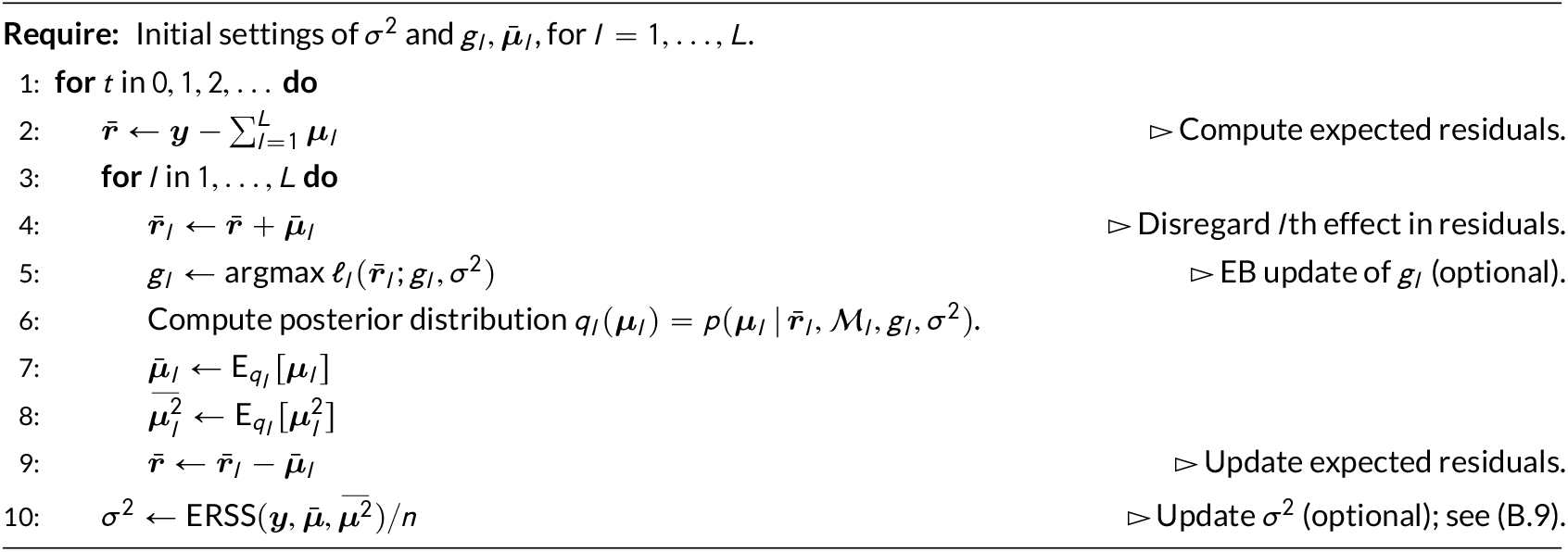

The update for *σ*^2^ in Algorithm A1 is easily obtained by taking partial derivative of (B.6), setting to zero, and solving for *σ* ^2^,giving

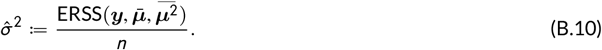

The update for *q_l_, g_l_* corresponds to finding the EB solution for the simpler (single effect) model 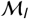 in which the data ***y*** are replaced with the expected residuals,

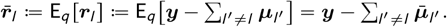

The proof of this result is given below in Proposition A1.

Substituting these ideas into Algorithm A1 yields Algorithm A2, which generalizes the IBSS algorithm (Algorithm 1) given in the main text.

### B.3 Special case of SuSiE model

The SuSiE model is a special case of the above additive effects model when ***μ***_*l*_ = ***Xb***_*l*_. In this case, 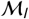 is the SER model, and the first and second moments of ***μ***_*l*_ are easily found from the first and second moments of ***b***_*l*_:

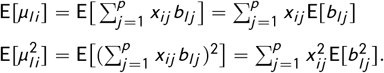

#### Algorithm A3 Iterative Bayesian stepwise selection (extended version)

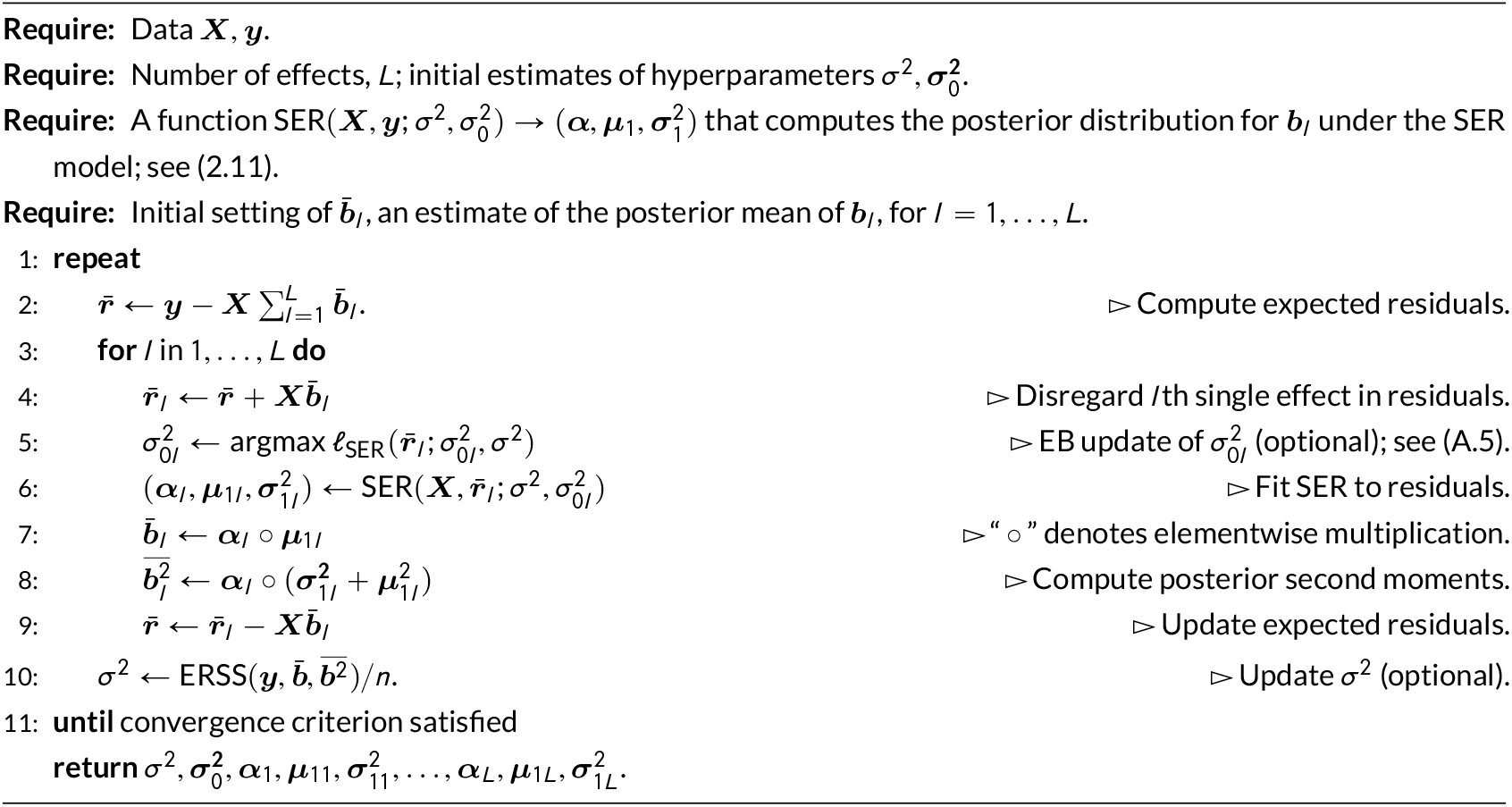

The expression for the second moment simplifies because only one element of ***b***
_*l*_ is non-zero under the SER model, and so 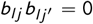 forany *j* ≠ *j*′. Becauseof this, we can easily formulate 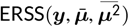 as a function of the first and second moments of ***b***_*l*_ — denoting this as 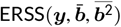 — and Algorithm A2 can be implemented using posterior distributions of ***b*** instead of posterior distributions of ***μ***.

For completeness, we give this algorithm, which is Algorithm A3. This algorithm is the same as the IBSS algorithm in the main text (Algorithm 1), with additional steps for fitting the hyperparameters *σ*^2^ and 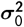. This is the algorithm implemented in the `susieR` software. The step to update 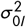 is a one-dimensional optimization problem; we implemented this step using the R function `optim`, which finds a stationary point of the likelihood surface with respect to 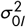. The algorithm terminates when the increase in the ELBO between successive iterations is smaller than a small non-negative number, *δ* (set to 0.001 unless otherwise stated). This is a commonly used stopping criterion in algorithms for fitting variational approximations.

### B.4 Update for *q_l_, g_l_* in additive effects model is EB solution for simpler model, 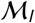

Here we establish that the update to *q_l_, g_l_* in Algorithm A1 can be implemented as the EB solution for 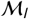 (Steps 5 and 6 in Algorithm A2). This result is formalized in the following proposition, which generalizes Proposition 1 in the main text.

#### Proposition A1.

*The q_l_, g_l_ that maximizes F in* (B.6), *the ELBO for the additive model*, 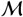, *can be found by maximizing the ELBO for the simpler model*, 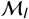, *in which the observed responses **y** are replaced by the expected residuals*, 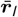:

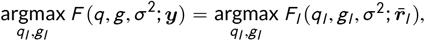

*where* 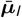 *is the vector of posterior mean effects defined above (see eq. B.7), and we define*

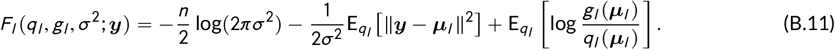

*Proof*. Omitting terms in the expression for *F* (from eq. B.6) that do not depend on *q_l_, g_l_* (these terms are captured by “const”), we have

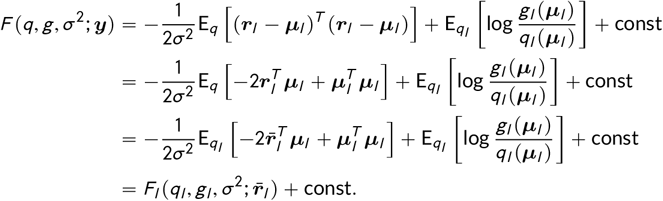

Further note that the optimization of *F_l_* does not restrict *q_l_*, so the maximum yields the exact EB solution for *M_l_* (refer to Section B.1.1); that is, 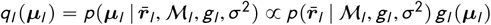 at the maximum.

### B.5 Convergence of IBSS algorithm

#### B.5.1 Proof of Corollary 1

##### Proof.

Step 5 of Algorithm 1 is simply computing the right-hand side of (3.9), in which the posterior distribution is determined by parameters 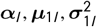. Therefore, by Proposition 1, it is a coordinate ascent step for optimizing the /th coordinate of 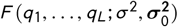 in which *q_l_* is determined by the parameters 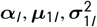.

#### B.5.2 Proof of Proposition 2

##### Proof.

By Proposition 2.7.1 of Bertsekas (1999), the sequence of iterates *q* converges to a stationary point of *F* provided that 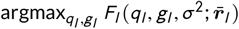 is uniquely attained for each *l*. When 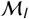 is the SER model and ***μ***_*l*_ = ***Xb***_*l*_, the lower bound *F_l_* (B.11) is

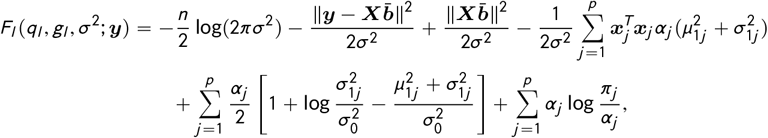

To lighten notation in the above expression, the *l* subscript was omitted from the quantities ***α*** = (*α*_1_,…, *α_p_*), ***μ***_1_ = (*μ*_11_,…, *μ*_1*p*_) and 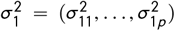 specifying the SER approximate posterior, *q_l_*, and likewise for the vector of posterior means, 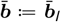 with elements 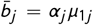. Taking partial derivatives of this expression with respect to the parameters ***α, μ***_1_ and 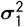, the maximum can be expressed as the solution to the following system of equations:

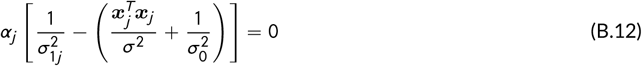

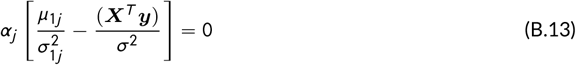

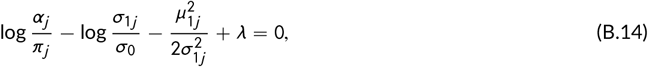

where 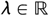 is an additional unknown, set so that *α*_1_ + ⋯ + *α_p_* = 1 is satisfied. The solution to this set of equations is finite and unique if 0 < *σ*, *σ*_0_ < ∞ and *π_j_* > 0 for all *j* = 1,…, *p*. Also note that the solution to (B.12–B.14) recovers the posterior expressions for the SER model.

### B.6 Computing the evidence lower bound

Although not strictly needed to implement Algorithms A2 and A3, it can be helpful to compute the objective function, *F* (e.g., to monitor the algorithm’s progress, or to compare solutions). Here we outline a practical approach to computing *F* for the SuSiE model.

Refer to the expression for the ELBO, *F*, given in (B.6). Computing the first term is straightforward. The second term is the ERSS (B.9). The third term can be computed from the marginal log-likelihoods *ℓ_l_* in (B.5), and computing this is straightforward for the SER model, involving a sum over the *p* possible single effects (see eq. A.5). This is shown by the following lemma:

#### Lemma A1.

*Let* 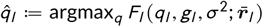. *Then*

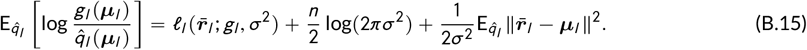

*Proof*. Rearranging (B.11), and replacing ***y*** with 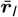, we have

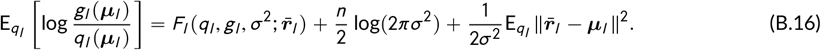

The result then follows from observing that *F_l_* is equal to *ℓ_l_* at the maximum, 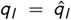; that is, 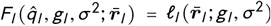.

### B.7 Expression for the expected residual sum of squares (ERSS)

The expression (B.9) is derived as follows:

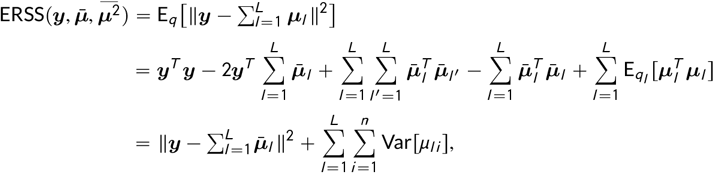

where 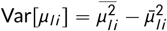.

## C CONNECTING SUSIE TO STANDARD BVSR

When *L* ≪ *p*, the SuSiE model (3.1–3.6) is closely related to a standard BVSR model in which a subset of *L* regression coefficients are randomly chosen to have non-zero effects.

To make this precise, consider the following regression model:

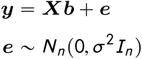

with *n* observations and *p* variables, so that ***b*** is a *p*-vector. Let 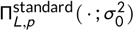 denote the prior distribution on ***b*** that first randomly selects a subset 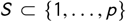 uniformly among all 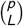 subsets of cardinality 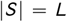, and then randomly samples the non-zero values 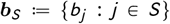 independently from 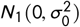, setting the other values 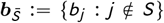 to 0. (This is a version of the prior considered by Castillo et al. 2015, with 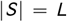.) Further, let 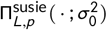 denote the prior distribution on ***b*** induced by the SuSiE model (3.1–3.6) with identical prior variances, 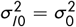, for all *l* = 1,…, *L*.

### Proposition A2.

*With L fixed, letting p* → ∞, *the SuSiE prior is equivalent to the standard prior. Specifically, for any event A*,

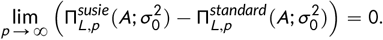

*Proof*. Fix *L* and *p*, and let *B* denote the event that the *L* vectors *γ*_1_,…, *γ_L_* in the SuSiE model are distinct from one another. Conditional on *B*, it is clear from symmetry that the SuSiE prior exactly matches the standard prior; that is, 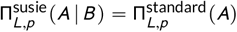, dropping notational dependence on 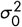 for simplicity. Thus,

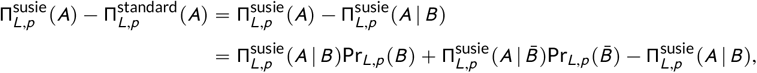

where the last line follows from the law of total probability. The result then follows from the fact that the probability Pr_*L,p*_(*B*) → 1 as *p* → ∞:

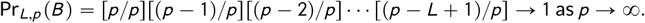

## D SIMULATION DETAILS

### D.1 Simulated data

For the numerical simulations of eQTL fine mapping in Section 4, we used *n* = 574 human genotypes collected as part of the Genotype-Tissue Expression (GTEx) project (GTEx Consortium, 2017). Specifically, we obtained genotype data from whole-genome sequencing, with imputed genotypes, under dbGaP accession `phs000424.v7.p2`. In our analyses, we only included SNPs with minor allele frequencies 1% or greater. All reported SNP base-pair positions were based on Genome Reference Consortium (GRC) human genome assembly 38.

To select SNPs nearby each gene, we considered two SNP selection schemes in our simulations: (i) in the first scheme, we included all SNPs within 1 Megabase (Mb) of the gene’s transcription start site (TSS); (ii) in the second, we used the *p* = 1,000 SNPs closest to the TSS. Since the GTEx data contain a very large number of SNPs, the 1,000 closest SNPs are never more than 0.4 Mb away from the TSS. Selection scheme (i) yields genotype matrices ***X*** with at least *p* = 3,022 SNPs and at most *p* = 11,999 SNPs, and an average of 7,217 SNPs.

For illustration, correlations among the SNPs for one of the data sets are shown in Fig. A1 (see also Fig. 1).

**FIGURE A1.**
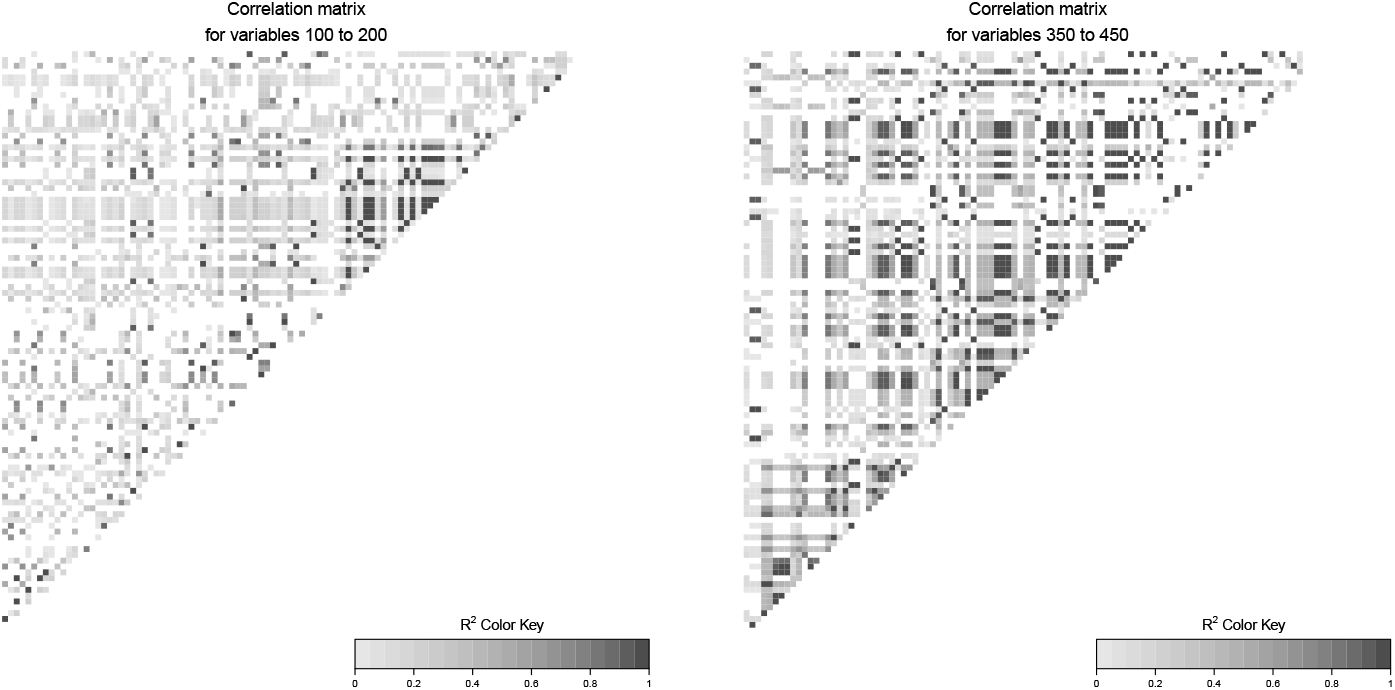
Correlations among variables (SNPs) in an example data set used in the fine mapping comparisons. Left-hand panel shows correlations among variables shown at positions 100–200 in Fig. 1; right-hand panel shows correlations among variables shown at positions 350–450. For more details on this example data set, see Section 4.1 in the main text.

### D.2 Software and hardware specifications for numerical comparisons study

In CAVIAR, we set all prior inclusion probabilities to 1 /*p* to match the default settings used in other methods. In CAVIAR and FINEMAP, we set the maximum number of effect variables to the value of 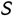 that was used to simulate the gene expression data. The maximum number of iterations in FINEMAP was set to 100,000 (this is the FINEMAP default). We estimate *σ*^2^ in SuSiE for all simulations.

All computations were performed on Linux systems with Intel Xeon E5-2680 v4 (2.40 GHz) processors. We ran SuSiE in R 3.5.1, with optimized matrix operations provided by the dynamically linked OpenBLAS libraries (version 0.3.5). DAP-G and CAVIAR were compiled from source using GCC 4.9.2, and pre-compiled binary executables, available from the author’s website, were used to run FINEMAP.

## E FUNCTIONAL ENRICHMENT OF SPLICE QTL FINE MAPPING

To strengthen results of Section 5, here we provide evidence that splice QTLs identified by SuSiE are enriched in functional genomic regions, thus likely to contain true causal effects. To perform this analysis, we labelled one CS at each intron the “primary CS.” We chose the CS with highest purity at each intron as the primary CS; any additional CSs at each intron were labelled as “secondary CSs.” We then tested both primary and secondary CSs for enrichment of biological annotations by comparing the SNPs inside these CSs (those with PIP > 0.2) against random “control” SNPs outside all primary and secondary CSs.

We tested for enrichment of the same generic variant annotations used in Li et al. (2016). These include LCL-specific histone marks (H3K27ac, H3K27me3, H3K36me3, H3K4me1, H3K4me2, H3K4me3, H3K79me2, H3K9ac, H3K9me3, H4K20me1), DNase I hypersensitive sites, transcriptional repressor CTCF binding sites, RNA polymerase II (PolII) binding sites, extended splice sites (defined as 5 base-pairs upstream and downstream of an intron start site, and 15 base-pairs upstream and downstream of an intron end site), as well as intron and coding annotations. In total, 16 variant annotations were tested for enrichment.

Figure A2 shows the enrichments in both primary and secondary CSs for the 12 out of 16 annotations that were significant at *p*-value < 10^−4^ in the primary signals (Fisher’s exact test, two-sided, no *p*-value adjustment for multiple comparisons). The strongest enrichment in both primary and secondary signals was for extended splice sites (odds ratio « 5 in primary signals), which is reassuring given that these results are for splice QTLs. Other significantly enriched annotations in primary signals include PolII binding, several histone marks, and coding regions. The only annotation showing a significant depletion was introns. Results for secondary signals were qualitatively similar to those for primary, though all enrichments are less significant, which is most likely explained by the much smaller number of secondary CSs.

**FIGURE A2.**
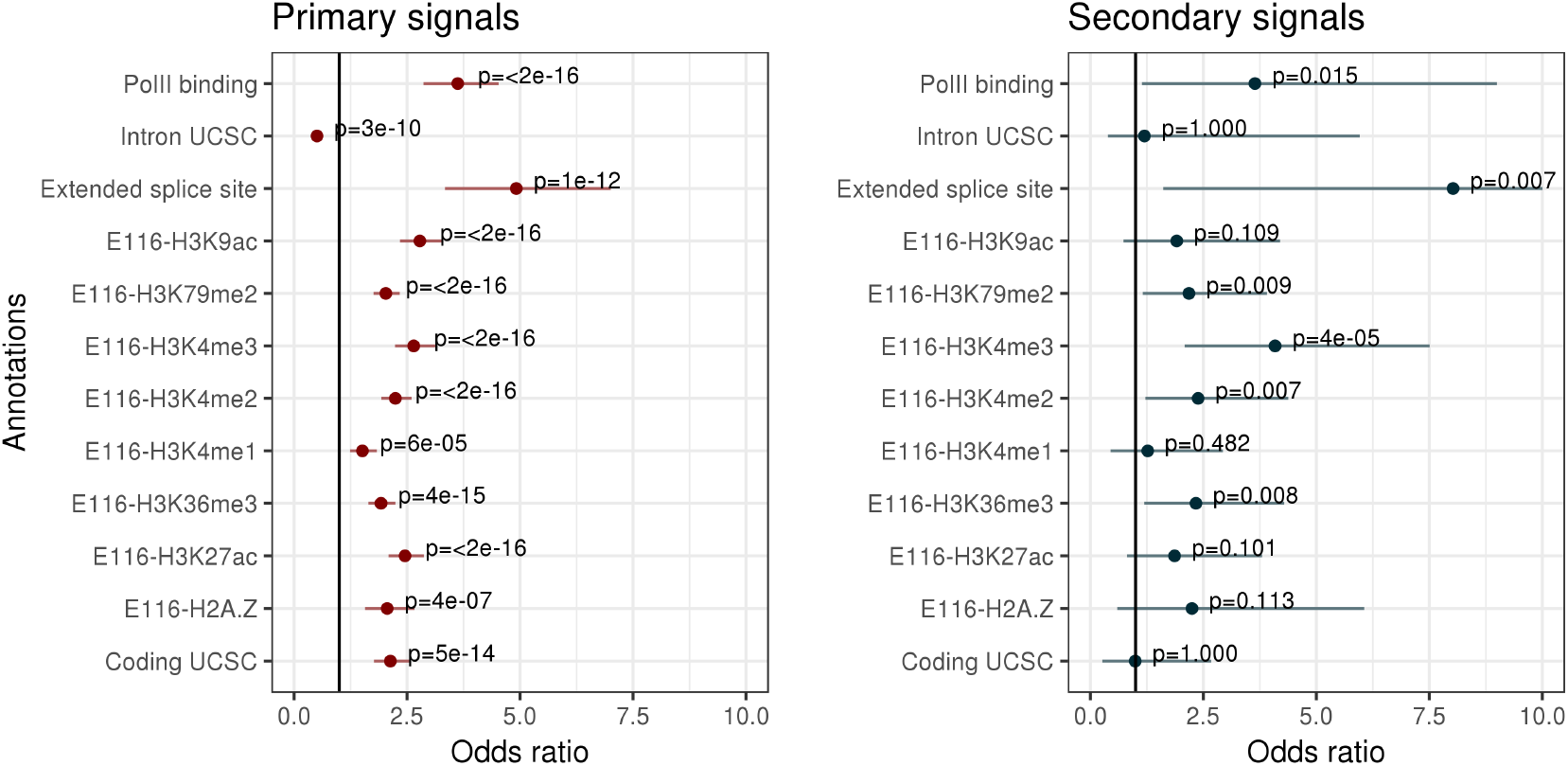
Splicing QTL enrichment analysis results. Estimated odds ratios, and + 2 standard errors, for each variant annotation, obtained by comparing the annotations of SNPs inside primary/secondary CSs against random “control” SNPs outside CSs. The *p*-values are from two-sided Fisher’s exact test, without multiple testing correction. The vertical line in each plot is posited at odds ratio = 1.

## Supplementary Figures

**FIGURE S1.**
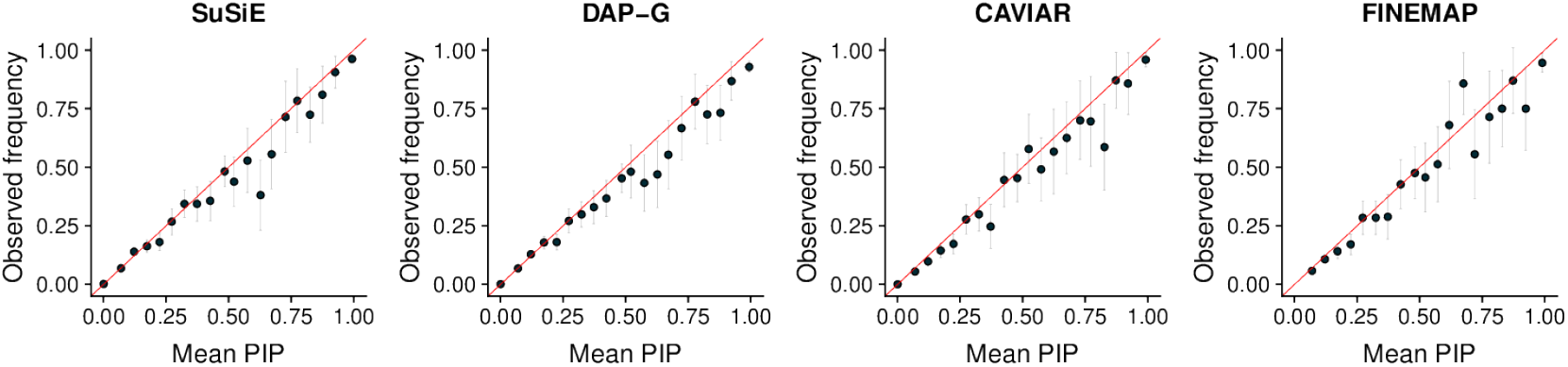
Assessment of PIP calibration. Variables across all simulations were grouped into bins according to their reported PIP (using 20 equally spaced bins, from 0 to 1). The plots show the average PIP for each bin against the proportion of effect variables in that bin. A well calibrated method should produce points near the *x = y* line (the diagonal red lines). Gray error bars show +2 standard errors.

**FIGURE S2.**
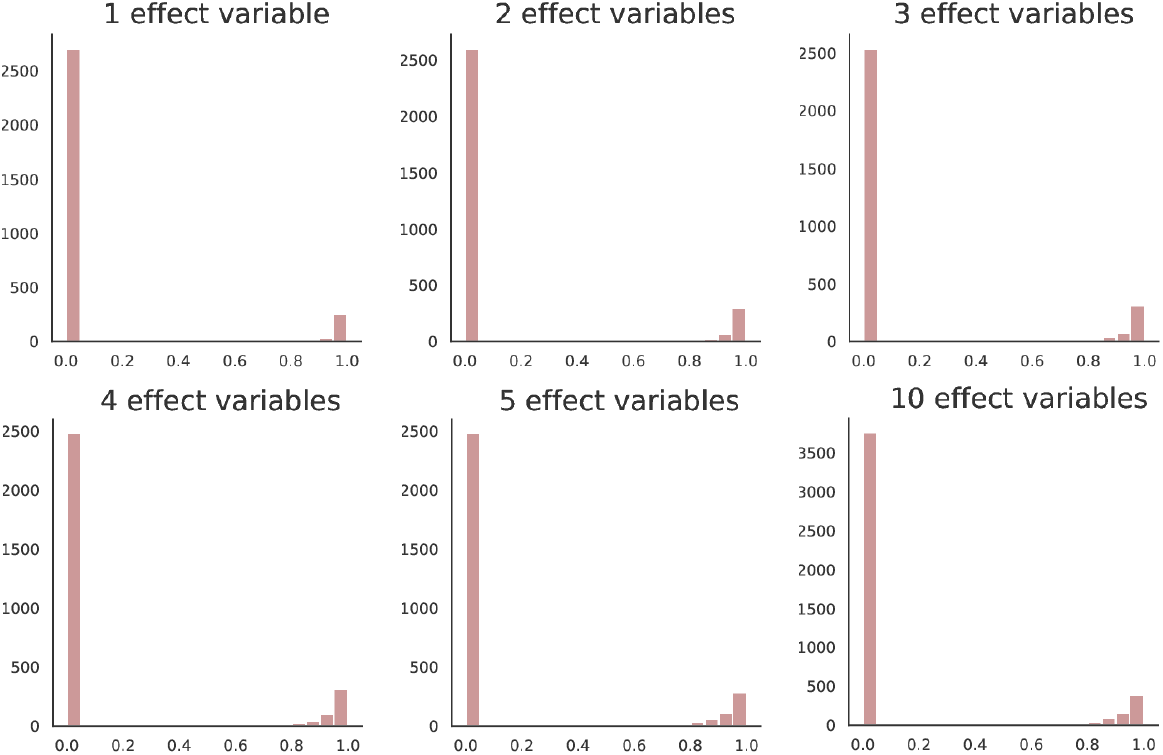
Distribution of purity for 95% credible sets for different numbers of effect variables. Histograms for 1–5 effect variables are obtained from all 95% credible sets produced by SuSiE in the first simulation scenario, with 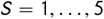, as described in Section 4 of the main text, and the 10 effect variables histogram is obtained from all 95% credible sets produced by SuSiE in the second simulation scenario, with 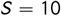.

**FIGURE S3.**
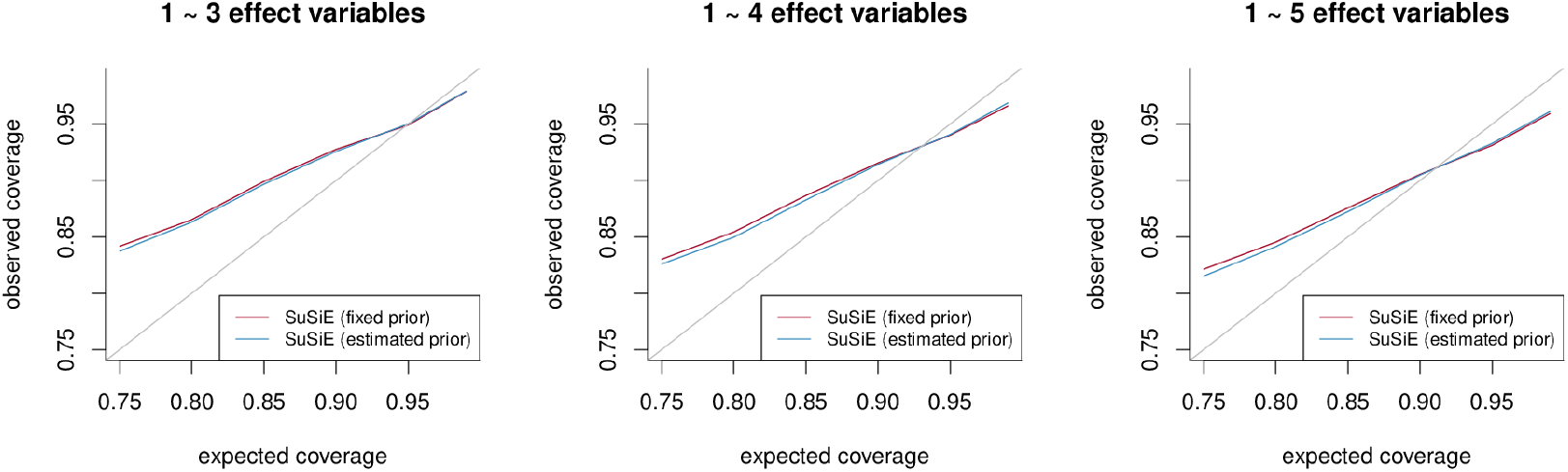
Additional assessment of SuSiE CS coverage. These three plots show coverage of SuSiE credible sets as *ρ* (the probability that the credible set contains at least one effect variable; see Definition 1 in the main text) is varied from 75% to 99%. Proportions shown in the vertical axis are based on all credible sets generated by SuSiE in simulations from simulation scenario 1, with different simulation settings for 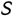, the number of effect variables. Consistent with Fig. 3, coverage decreases with the inclusion of weaker signals.

**FIGURE S4.**
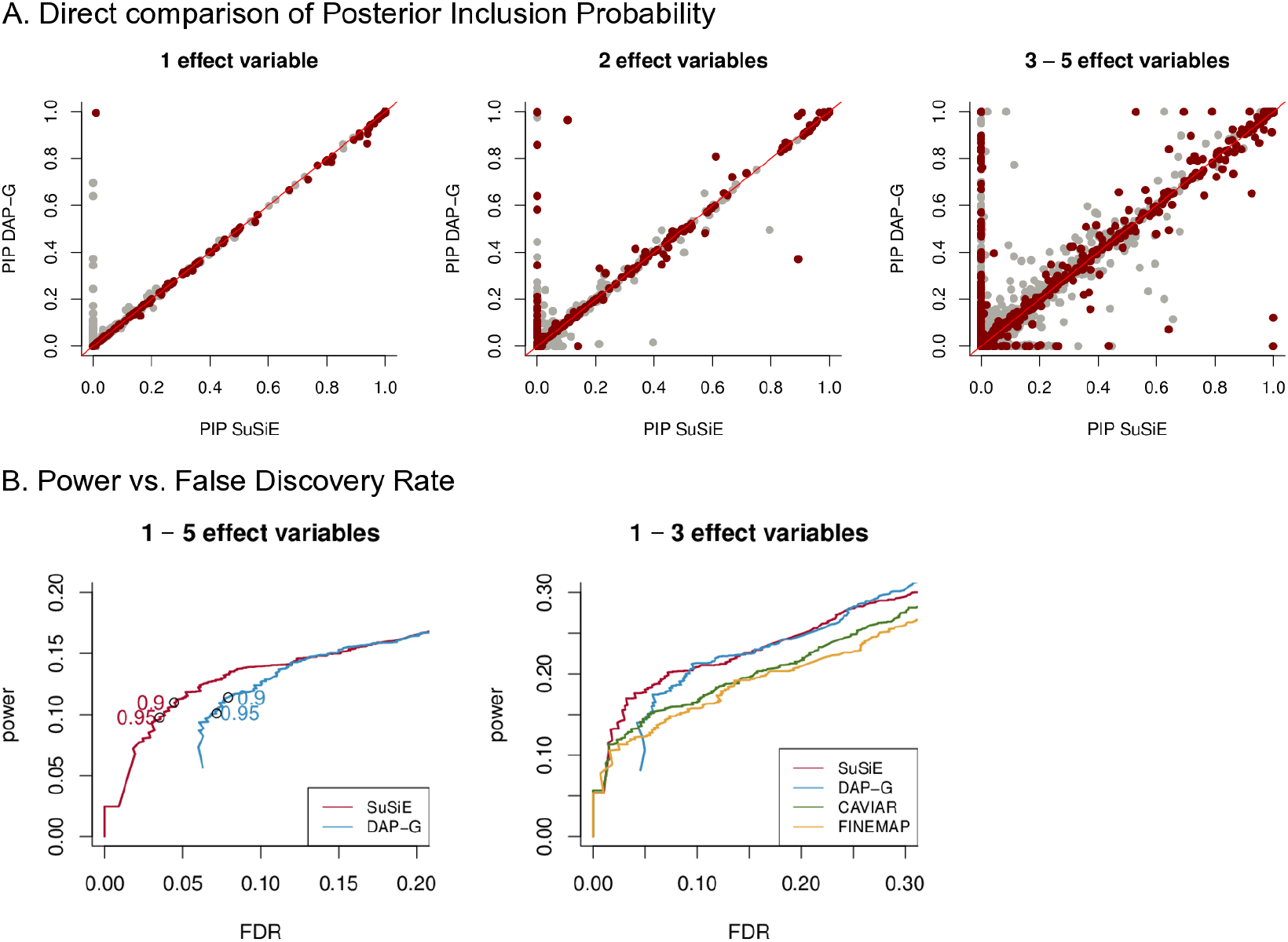
Comparison of posterior inclusion probabilities (PIPs) computed by SuSiE, in which the prior variances ***σ***^2^ are estimated rather than fixed to 0.1, against PIPs computed by DAP-G, and by other methods. The results shown here from methods other than SuSiE are the same as the results in Fig. 2. For an explanation of the individual plots, see Fig. 2.

**FIGURE S5.**
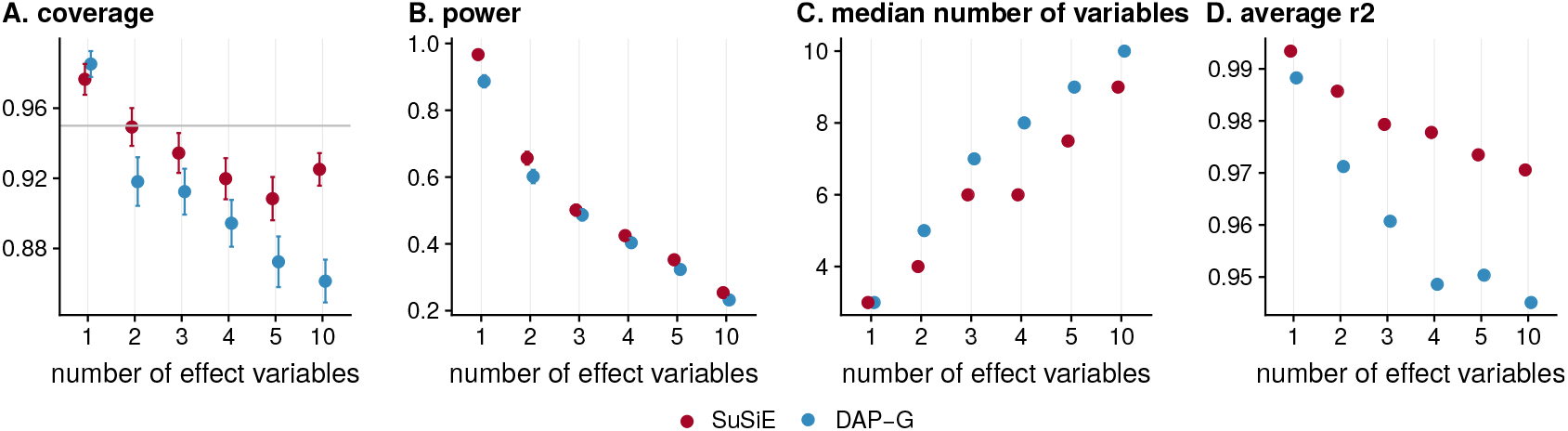
Comparison of 95% credible sets (CS) from SuSiE, in which the prior variances ***σ***^2^ are estimated rather than fixed to 0.1, and DAP-G: (A) coverage, (B) power, (C) median size, and (D) average squared correlation among variables in each credible set. The DAP-G results shown here are the same as the DAP-G results shown in Fig. 3. For an explanation of the individual plots, see Fig. 3.

